# Feasibility of Precision Functional Mapping in Youth Multi-Echo fMRI Data

**DOI:** 10.64898/2026.05.20.726578

**Authors:** Isaac N. Treves, David Pagliaccio, Gaurav H. Patel, Reem Tamimi, Jihoon A. Kimerty, Randy P. Auerbach, Hilary A. Marusak

**Author notes:** Corresponding Author, 2412 Herbert Pardes Building, New York State Psychiatric Institute, New York, NY 10032, USA.

## Abstract

There is growing interest in identifying brain function underlying adolescent cognition, personality, and psychopathology. One promising approach is Precision Functional Mapping (PFM) of MRI functional connectivity, a data-intensive method for characterizing individualized brain networks. Foundational studies suggest that PFM can detect stable, task-responsive, and clinically relevant networks. Studies demonstrate that both functional connectivity reliability and network stability improve with increasing data quantity, although benchmark estimates vary across populations, preprocessing pipelines, and MRI acquisition approaches. Accordingly, it is important to understand how PFM performs in adolescent populations and with multi-echo fMRI acquisition. In a case study of eight youth (ages 10-17), we applied PFM to 80-minutes of combined resting-state and task-based fMRI. The resulting networks were highly modular, consistent with adult templates, and without evidence of structural registration artifacts. Functional connectivity reliability compared favorably to prior single-echo studies, with multivariate similarity and ICC estimates showing early stabilization around 10-15 minutes despite continued improvement with additional data. Trait-like stability increased gradually with acquisition time and a Bayesian algorithm (MS-HBM) demonstrated higher stability than Infomap. Across algorithms, stability was greatest in sensory networks (somatomotor, auditory, visual). Furthermore, when evaluating task-based responses to threat and attention paradigms, only the auditory network consistently benefited from individualized mapping over group template networks. These findings suggest that, with constrained scanning time, PFM is especially effective for characterizing sensory and perceptual networks in adolescents. Bridging the methodological divide between deeply sampled individual cases and large-scale developmental studies will require further innovation and validation.

## Introduction

The identification of stable, individualized brain networks is of considerable interest in functional magnetic resonance imaging (fMRI). Though much research has relied on standardized, group-average template networks, individualized networks may better predict treatment responses, capture heterogeneity in psychiatric disorders, and optimize neuromodulation approaches for a given patient (Gratton et al., 2020; Labonte et al., 2024; Lee et al., 2024; Lynch et al., 2024). Precision functional mapping (PFM) involves estimating networks based on functional connectivity (FC) relationships across the cortex or entire brain and has emerged as a promising framework for this purpose. Foundational PFM studies often leveraged densely sampled data (> 2 hours of resting-state fMRI from individual participants across sessions; Demeter et al., 2025; Gordon, Laumann, Gilmore, et al., 2017; Laumann et al., 2015) to establish that individual-specific network organization is reproducible within a person and includes idiosyncratic features not apparent in group-average maps (Gordon, Laumann, Gilmore, et al., 2017; Gordon et al., 2020; Gratton et al., 2020; Seitzman et al., 2019). Studies also indicate that data needs are target dependent: whole connectome FC estimates can converge relatively quickly (∼30 minutes in adults; Gordon et al., 2017, Laumann et al., 2015), while reliable network assignment and topographic stability can require much longer duration (∼90 minutes; Gordon et al., 2017, Laumann et al., 2015). This variability raises a central translational question: what level of individualized network characterization is achievable under the more constrained scanning conditions typical of developmental and clinical research?

Adult PFM studies have utilized a variety of methods for identifying individualized networks. Methods include Infomap (Rosvall & Bergstrom, 2008), Homologous Functional Region mapping (Wang et al., 2015), Template matching (Seitzman et al., 2019), and multi-session hierarchical Bayesian model (MS-HBM (Kong et al., 2019). Original research on adult PFM relied on Infomap, a computationally intensive method that derives whole-brain networks (including subcortex) by subdividing communities of vertices or voxels based on functional connectivity thresholds. Individual networks derived from Infomap often differed topologically (in spatial location) from group-average networks (Laumann et al., 2015), and the stability of these differences over time has been a central focus. Stability analyses suggest that individualized network topologies become more stable with additional data, reaching elbows (points of diminishing returns) at around 60 minutes (Gordon, Laumann, Gilmore, et al., 2017). In addition, network topologies appear similar across resting-state or task-based MRI, allowing data concatenation to increase usable signal (Gratton et al., 2018; Wang et al., 2015).

Stable networks may be identified across the brain, though they are most consistently observed in primary sensory cortices (Gordon, Laumann, Adeyemo, et al., 2017; Hakonen et al., 2025; Moser et al., 2024)—including visual, somatomotor, and auditory networks. Interestingly, these areas also show stronger task-related activations when individualized networks are used instead of group templates, suggesting they are meaningfully related to individual perceptual abilities (Gordon, Laumann, Gilmore, et al., 2017; Laumann et al., 2015). In contrast, regions susceptible to signal dropout (e.g., ventromedial prefrontal cortex, anterior medial temporal lobe), exhibit more limited stability (Demeter et al., 2025; Rai et al., 2025).

A related concept to network topological stability is reliability of FC. FC reflects bidirectional correlations between brain regions (Biswal et al., 1995) and is commonly used as a putative biomarker of cognition, psychopathology, and development (Gabrieli et al., 2015). As FC measures are utilized in precision mapping algorithms, the reliability of FC relates to the stability of individualized networks. However, FC reliability is often modest, influenced by head motion, task demands, and acquisition length (Finn, 2021; Noble et al., 2017, 2019). Adult PFM studies show that FC reliability improves with data quantity: multivariate similarity grows rapidly reaching elbows at ∼30 minutes of high-quality data (Gordon, Laumann, Gilmore, et al., 2017), though other studies suggest longer acquisitions (e.g., ∼90 minutes (Raut et al., 2019).

Alternative acquisition techniques may alter reliability. Multi-echo fMRI collects signal at multiple points (“echoes”) following the RF pulse (Kundu et al., 2017). Given that blood-oxygenation responses and artifacts follow distinct decay curves, multi-echo allows for model-driven denoising, in contrast to the heuristic filtering approaches applied in conventional single-echo fMRI (DuPre et al., 2021). Studies consistently show that multi-echo improves signal-to-noise ratio relative to single-echo (Kundu et al., 2017). Accordingly, researchers have begun employing multi-echo with PFM, with early evidence in adults suggesting it may reduce the scan time required for stable FC and network topological stability (Lynch et al., 2020).

### Studies in Youth

PFM in youth offers a powerful approach to study neuromaturation and the development of psychopathology (Labonte et al., 2024). One study in children (ages 8-11) found that 45-50 minutes of resting-state was sufficient for high functional connectivity reliability (multivariate similarity; Demeter et al., 2025). They also found that children exhibited lower between-person variability than adults, suggesting that development leads to more network differentiation. A study of multi-echo fMRI in children (ages 6-9) passively viewing videos reported lower acquisition-time estimates for low-motion participants (∼24 minutes to reach high similarity), although edge-wise ICC estimates generally did not reach fair reliability until approximately 50 minutes for most networks (Rai et al., 2025). Independent effects on reliability were found for age, head motion, and task condition. These studies primarily focused on functional connectivity reliability and/or group comparisons to adults (Demeter et al., 2025), and less is known about the spatial stability of individualized networks across acquisition durations, particularly in adolescent brain data.

The present study builds on this prior work by examining both reliability (integrating multiple methods including ICC) and spatial stability in youth ages 10–17 using multi-echo fMRI. We extend prior pediatric PFM work by evaluating additional individualized network estimation approaches like Infomap and MS-HBM. Further, we systematically examine the topological (spatial) stability of individualized networks and their task responsiveness. In doing so, we contribute to emerging findings that sensory and higher-order networks may have different stabilities (Gordon, Laumann, Gilmore, et al., 2017; Hakonen et al., 2025; Moser et al., 2024). Typically, in adults, case studies have proved an effective way to address these questions about PFM, as constrained samples allow for: (a) careful quality assurance, (b) repeated, intensive computational methods, and (c) in-depth visualization. Accordingly, the current study provides a preregistered, systematic evaluation of multi-echo PFM in a case study of 8 youth (ages 10-17) using a feasible 80-minute protocol. To maximize available data, we chose to include all data across task and resting-state conditions (“mixed-state data”)(Hermosillo et al., 2024; Wang et al., 2015), while conducting sensitivity analyses to understand task condition effects.

## Methods

### Preregistration

A preregistered analysis plan may be found at https://osf.io/8rb4n. In the preregistration, the patch mapping pipeline was discussed (Wang et al., 2015). Patch mapping performance was suboptimal in terms of network coverage, and so we instead used a hierarchical Bayesian method, MS-HBM, which has been previously compared to Infomap (Lynch et al., 2024). Additionally, we did not characterize network variants (e.g., ectopic intrusions), as this study focused on the statistical properties of the maps.

### Data

Multi-echo fMRI and phenotypic data were collected as part of a neuroimaging study conducted in a community sample of children and adolescents (10-17 years old; Marusak et al., 2024). Inclusion criteria consisted of age range 10-17 and right-handedness. Exclusion criteria consisted of lifetime autism or developmental disorders, bipolar disorder, psychotic disorder, obsessive-compulsive disorder, pregnancy, neurological disorders, significant head injury with ongoing symptoms, substance use at the time of scanning (biologically verified), oral contraceptive use, and non-stimulant psychotropic medication use. Data were collected at Wayne State University on a Siemens Verio 3T system with a 32-channel head coil (full parameters found in **Supplement Text 1**). Participants contributed up to 80 minutes of data over a single or two consecutive days, including four eyes-closed resting-state runs and five task-based fMRI paradigms: fear conditioning, fear extinction, extinction recall, fear renewal (Marusak et al., 2017, 2018), and the Attention Network Task (Fan et al., 2005) (The run order is provided in **Supplement Text 2**). For the current case study, eight participants were selected based on relatively low head motion and adequate task performance (**Supplement Text 3)**. The eight individuals were selected to be diverse in age, race, and biological sex (**Table 1**). All procedures complied with the Declaration of Helsinki and were approved by the Wayne State University Institutional Review Board. A parent or legal guardian provided written informed consent, and all youth provided informed assent.

**Table 1.**
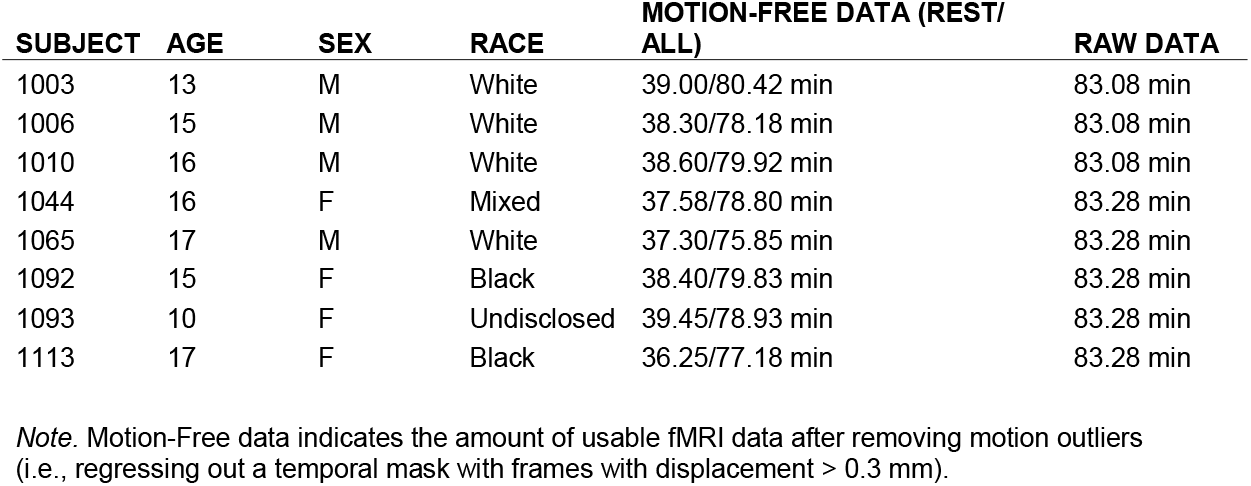
Participant information.

| SUBJECT | AGE | SEX | RACE | MOTION-FREE DATA (REST/<br>ALL) | RAW DATA |
| --- | --- | --- | --- | --- | --- |
| 1003 | 13 | M | White | 39.00/80.42 min | 83.08 min |
| 1006 | 15 | M | White | 38.30/78.18 min | 83.08 min |
| 1010 | 16 | M | White | 38.60/79.92 min | 83.08 min |
| 1044 | 16 | F | Mixed | 37.58/78.80 min | 83.28 min |
| 1065 | 17 | M | White | 37.30/75.85 min | 83.28 min |
| 1092 | 15 | F | Black | 38.40/79.83 min | 83.28 min |
| 1093 | 10 | F | Undisclosed | 39.45/78.93 min | 83.28 min |
| 1113 | 17 | F | Black | 36.25/77.18 min | 83.28 min |

*Note.* Motion-Free data indicates the amount of usable fMRI data after removing motion outliers (i.e., regressing out a temporal mask with frames with displacement > 0.3 mm).

### Data preprocessing and denoising

Preprocessing, denoising, and quality assurance steps were applied to fMRI data using identical multi-echo processing pipelines detailed in Lynch et al., 2024. As fieldmaps were not collected, we deployed an average fieldmap template (Treiber et al., 2016) (**Supplement Text 4**). The preprocessing pipeline includes, briefly: fieldmap correction, discarding 10 volumes (to allow for steady-state), co-registration of average steady-state images to ACPC-aligned structural images, global mean time series regression, and brain extraction. Structural data were preprocessed using the HCP PreFreesurfer, FreeSurfer, and PostFreeSurfer pipelines (Glasser et al., 2013). Data were analyzed in CIFTI format consisting of volumetric (subcortex) and surface (cortex) data structures.

For further multi-echo preprocessing and denoising, *tedana* was used (DuPre et al., 2021), retaining BOLD components (showing T2 signal decay) and discarding noise components (S0-dependent) (Kundu et al., 2013). Per previous work, we only selected components showing a correlation with group networks > 0.1. Head motion correction consisted of regressing out a temporal mask which removed frames with > 0.3 mm of framewise displacement (FD) (Lynch et al., 2024). Signal complexity (number of components) and head motion (FD) were compared to individualized network sizes to assess sensitivity.

For a pseudo single-echo comparison, the middle-echo images were isolated (Lynch et al., 2020), and then preprocessed identically. Instead of *tedana* denoising, the ICA-AROMA denoising method was applied (Lynch et al., 2024; Pruim et al., 2015).

### Individualized Networks

For whole-brain networks (including subcortex, brainstem, and cerebellum), the Infomap algorithm was applied for community detection on sparse graphs of FC (Gordon, Laumann, Gilmore, et al., 2017; Lynch et al., 2024; Rosvall & Bergstrom, 2008). Infomap was conducted with a smoothing kernel equivalent to 6mm FWHM and a graph density (sparse threshold) of 0.1% (Lynch et al., 2024), which has previously been shown to maximize homogeneity and task responsiveness (Gordon et al., 2020). In the current study, Infomap resulted in a mean of 92.75 communities per individual (range 74–108). Each vertex or voxel was assigned to a single community, and the communities were algorithmically assigned to one of 20 possible functional network identities primarily according to their FC and spatial locations relative to a specified set of priors (Gordon et al., 2016). These networks included: Default-Parietal, Default-Anterolateral, Default-Dorsolateral, Default-Retrosplenial, Visual-Lateral, Visual-Stream, Visual-V1, Visual-V5, Frontoparietal, Dorsal Attention, Premotor/Dorsal Attention II, Language, Salience, Cingulo-opercular/Action-mode, Parietal memory, Auditory, Somatomotor-Hand, Somatomotor-Face, Somatomotor-Foot, and Auditory or Somato-Cognitive-Action. Low-confidence assignments were reviewed manually. After identifying the networks, each network surface area was computed for each individual.

In an exploratory analysis, the stability of networks was examined using a hierarchical Bayesian model with a smoothing kernel equivalent to 4mm FWHM (Kong et al., 2019), which employs group connectivity priors to identify cortical networks (**Supplement Text 5)**.

### Reliability and Stability Measures

Approximately 80 minutes of cleaned mixed-state data (75-81 minutes) incorporating tasks and resting-state runs were supplied for each individual. Iterative 5% splits of the data were conducted (Gordon, Laumann, Gilmore, et al., 2017) to test the stability and reliability of various measures. A split-half analysis was conducted using 40 minutes of data as the “reference” (with each individual’s run order randomized) and incrementally larger portions of the remaining 40 minutes as the ‘test’. This split-half approach closely follows prior PFM validation methods (Gordon, Laumann, Gilmore, et al., 2017), where observed similarity is attributable to trait effects. To better understand the implications of mixed-state analyses, we conducted sensitivity analyses. In one analysis, we compared iterative splits for resting-state alone vs mixed-state. If stability or reliability increases faster across time for resting-state, this would indicate influence of task condition. To confirm these iterative results with the main split-half approach, we compared participants who had similar amounts of resting-state data across halves (‘balanced’) vs those who were unbalanced. This was conducted because independent splits within the resting-state data alone would entail significant restriction of data quantities.

Network stability analyses were applied using Normalized Mutual Information (NMI) for the aggregate networks (often used to compare categorical grouping variables) and Dice coefficients for each individual network (often used to determine the overlap between binary variables). Each time increment was separately run through Infomap and then compared using these measures. A high NMI or Dice represents high overlap (1 = perfect overlap). Elbow points in the stability curves were also assessed visually, as indicated by diminishing returns for additional data.

FC reliability was assessed by applying the communities derived from the entirety of data and calculating their *Z*-scored FC for randomly shuffled splits, with the shuffles incremented by 5% for each data point. We then derived measures of similarity (i.e., Pearson correlation between FC matrices) as well as graph-theoretical measures for the networks. Elbows were assessed visually, and confirmations were made using the Unit Invariant Knee approach, which identifies the point on a curve that is the maximum perpendicular distance from a line between the first and last points of the curve (Christopoulos, 2016; 2014). Elbows do not necessarily reflect asymptotes; instead, they reflect a change in the slope of improvement or points of diminishing returns, though reliability may continue to improve slowly with more data past the elbow.

Finally, we additionally examined homogeneity of time courses across voxels/vertices from each network, as measured by the variance explained by the first principal component wherein 0 = completely inhomogeneous and 100 = completely homogeneous. Given the lack of well-established findings on individualized network homogeneity, even in adults, these results are reported in the supplement. Briefly, results indicated little to no change with increased scan time, nor any improvements with regard to group networks (**Figure S1, S2**).

### Structural Confounds

We calculated vertex-level cortical thickness, registration distortion, and registration expansion using HCP processing files. These data were then averaged within each community from the Infomap pipeline. To estimate relationships with community size, we correlated community size and the average measures. Communities were chosen to provide more granularity than the networks, which are composed of sets of communities. A subsample of communities belonging to the frontoparietal network and salience network were examined for significant relationships (as frontal areas may be more affected by signal loss and subsequent registration effects).

### Task Responses

Task-based analyses were performed to examine whether individualized networks based on FC capture additional individual variation in task activation. Two tasks—the fear conditioning and Attention Network Task—were selected for activation contrast analysis. During the fear conditioning task (Marusak et al., 2017, 2018), aversive white noise stimuli were presented as the unconditioned stimulus (US). To model robust US-associated effects, we created an ‘auditory shock contrast’ consisting of the difference between white noise (US) and implicit baseline using an event-based general linear model (GLM) design with durations of zero for white noise. The Attention Network Task was modelled with three contrasts corresponding to alerting (center cue vs no cue), orienting (spatial vs center cue), and executive function (congruent or incongruent flanker), consistent with prior literature (Fan et al., 2005). Again, an event-based GLM with durations of zero was used.

Task responses were estimated using the *fitlins* 0.10.1 package, a BIDS-compatible linear modelling package (https://github.com/poldracklab/fitlins). Z-scored activation maps were then mapped to the surface, and cortical overlaps between the networks and the activation maps were assessed. Overlaps were displayed visually for particular examples and then, compared statistically to random shuffled versions of the networks as well as group template networks. Non-parametric tests were used to compare *Z*-scores averaged within random networks versus individualized networks, and then the non-parametric *p*-values were combined using Stouffer’s method to evaluate a *t*-statistic for the entire group of participants. To compare group networks and individualized networks, a one-group *t*-test between the difference in *Z*-scores and zero was conducted. Positive *t*-stats indicate greater activation for the individualized networks.

## Results

### Network Topology

Networks derived from multi-echo processing are shown in two exemplar participants in **Figure 1**. Networks showed a general correspondence with template networks (**Figure 2)**, and no significant correlations across all 8 participants with signal complexity (number of retained components from multi-echo denoising) nor head motion (FDR-*p*s > 0.8). Nevertheless, some modest descriptive differences from the group template (defined in adult populations) were observed. In particular, the default mode networks and visual lateral and streams networks appeared larger in our sample, whereas the language and default retrosplenial networks were smaller. When examining resting-state data only, network sizes were generally similar compared to the full data (**Figure S3**), although the salience network was larger on average.

Network quality in our single-echo comparison condition appeared inadequate (for both Infomap and MS-HBM). Single-echo derivatives tended to emphasize coarse gradients over network-specific motifs (**Supplement Text 7, Figure S4 & S5**).

**Figure 1.**
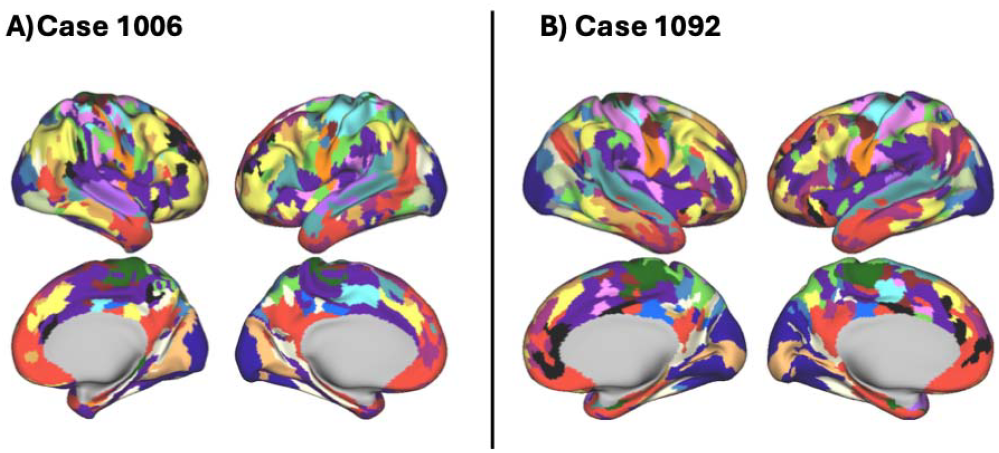
Example multi-echo networks.A) Case 1006. B) Case 1092. Colors represent each of 21 networks from Gordon templates and labels in Figure 2. Medial wall is gray.

**Figure 2.**
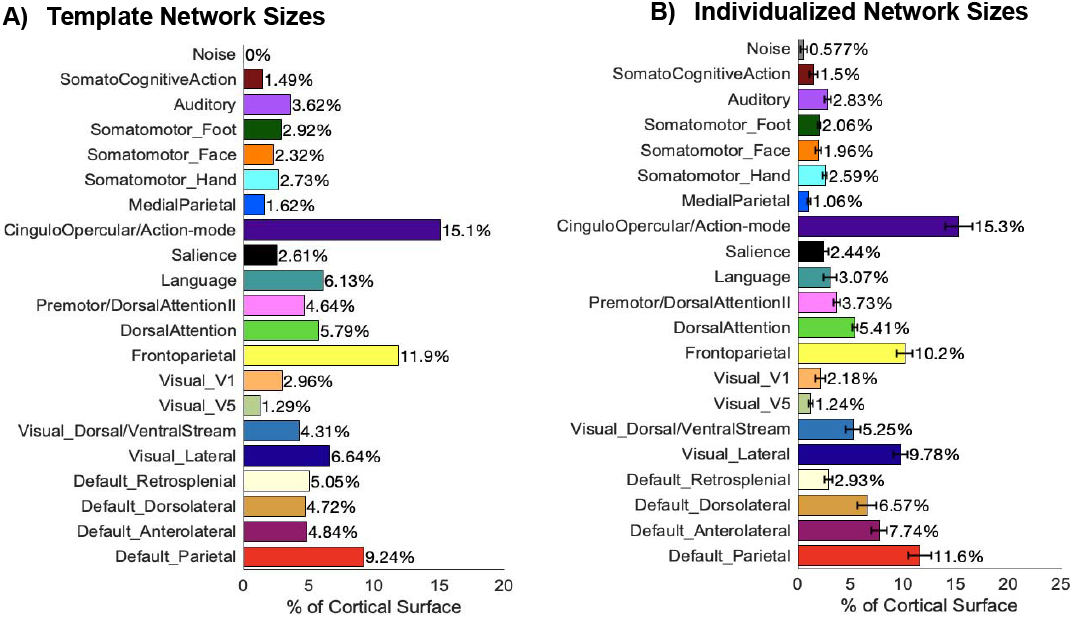
Comparison of Template Network Sizes and Individualized Network Sizes. In panel A), surface areas are shown for the cortical representations of the 20 Gordon template networks (+ Noise), based on adults. In panel B), the mean network template surface areas for the 8 participants (ages 10-17) reported here are shown, along with standard errors. The Noise network refers to communities from Infomap that were classified as Noise.

### Network Topology Stability

The stability of the networks over time was assessed using Dice coefficients for each network, and the NMI for all networks aggregated. In **Figure 3**, NMI is shown for the preregistered Infomap pipeline and the exploratory MS-HBM pipeline. For Infomap, the split-half overlap between networks increased only slowly with increasing amounts of data included, about 20% in 40 minutes, and this was not affected by altered parcellations (e.g., combining Default-Mode networks; **Figure S6**). For the Bayesian method, overlap was higher and appeared to grow faster, about 26% in 40 minutes. Split-half NMI did not reach the preregistered threshold of 0.75 for either pipeline. In addition, resting-state trajectories showed more rapid increases in stability than full mixed-state data trajectories (**Figure S7 & S8**).

**Figure 3.**
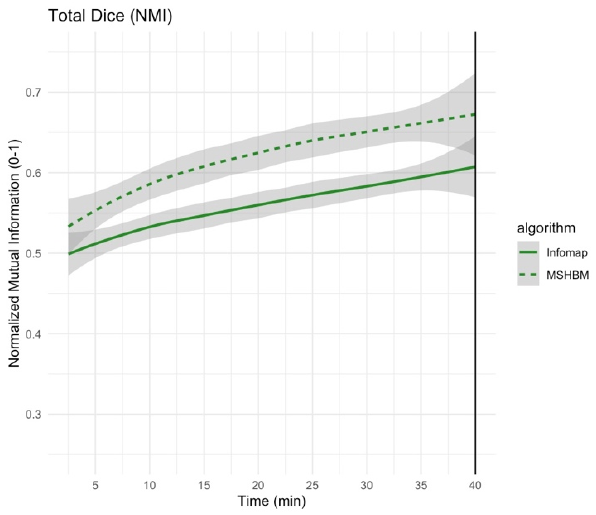
Normalized mutual information (similar to Dice) comparing the similarity of network topology across all networks over increasing time. In green, networks derived from the Infomap pipeline. In dashed green, networks derived from MS-HBM. Data were assessed in 2.5 minute (5%) increments, and interpolated between those points.

We also examined the differential stability of different networks (**Figure 4**). Somatomotor, auditory, and parietal networks demonstrate the highest dice coefficients compared to the independent split—about 2-3 times larger than salience and language networks. This distinction was consistent when examining the full mixed-state data or resting-state data only, and when examining MS-HBM networks (**Figure S9**). DMN subnetworks (parietal, anterolateral, dorsolateral, retrosplenial) exhibited moderate stability, but when combining them into one network the stability was the highest of any network (**Figure S10**). For Infomap, no visual elbows were identified, suggesting that all networks benefited from more data.

**Figure 4.**
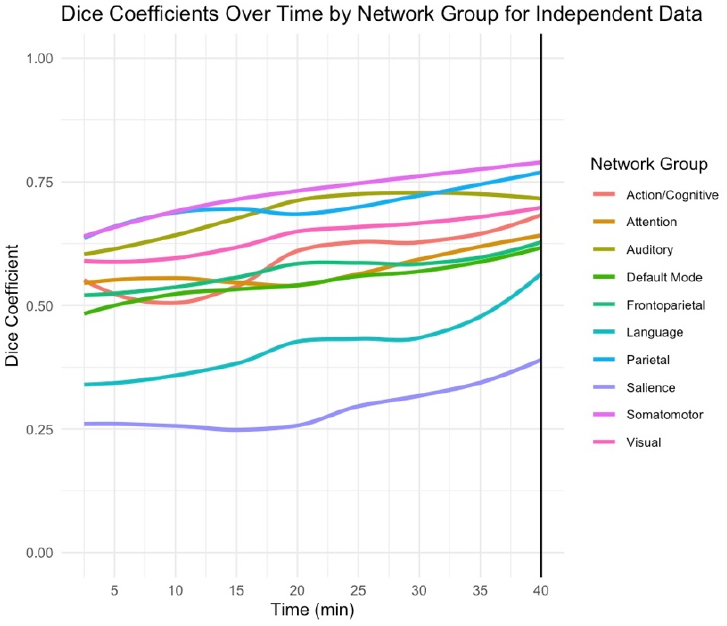
Dice coefficients for the Infomap networks over increasing minutes of the split, independent data. Networks are averaged into groups for viewing purposes. All visual networks are labelled Visual, all somatomotor networks are labelled Somatomotor, Action/Cognitive consists of SomatoCognitive Action and Cingulo Opercular, Dorsal Attention 1 and 2 are labelled ‘Attention’, and all default mode networks are labelled ‘Default Mode’. All other labels encompass unitary networks.

We also assessed vertex-wise stability by examining the consistency of network identification over data splits (**Figure S11, S12)**. Regions that showed high stability consisted of sensory areas like V1, superior temporal gyrus, and somatomotor and premotor areas. Interestingly, the ventromedial prefrontal cortex showed higher stability than adjoining frontal regions, perhaps because signal loss in that area resulted in more reliance on group templates. Regions that showed low stability consisted of dorsolateral PFC, ACC, anterior medial temporal lobes and posterior cingulate (likely influenced by instability in assignment of the DMN subnetworks).

### Network Connectivity

An example FC plot is found in **Figure S13**. The identified networks all show modularity (**Figure S14**). In addition, FC multivariate similarity was high (> 0.8) and reached elbows between 10-15 minutes of data (*M* = 12.19 min) (**Figure 5)**. FC similarity was highest when examining group template networks and lowest for vertices (**Figure S15**). Elbows for FC similarity and other graph-theoretic measures (**Figure S16**) were all found between 10-15 minutes. ICCs (intraclass correlation coefficients) also reached an elbow between 10-15 minutes (**Figure S17**), but exhibited sustained increases with more data. FC reliability did not seem systematically affected by resting-state vs mixed-state data (**Figure S18 & S19**).

**Figure 5.**
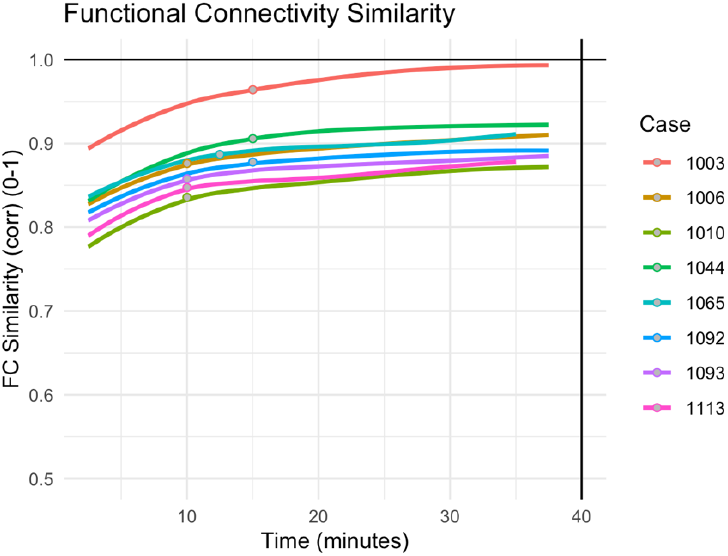
Functional connectivity (FC) similarity as function of the quantity of independent data, shown for each participant. The similarity was calculated as the correlation between FC in the subsample of independent data vs FC in the held-out data (40 minutes). Similarity was calculated for each 5% increment of the independent split-half data (up until 40 minutes). Elbows (points) may be observed at approximately 10-15 minutes of independent data, using the Unit Invariant Knee approach. For a discussion of Case 1003, which was characterized by extremely low motion, see **Supplement Text 6**.

### Structural Confounds

No significant correlations were found between network surface area and cortical thickness, registration expansion and contraction, nor registration strain (rotational changes) (*ps* > 0.3; **Figure S20**). When specifically examining frontal networks, like the frontoparietal network and salience network, no significant correlations were observed (*ps >* 0.25).

### Task Responses

Overlap between the activation *Z*-score maps and the individualized networks was assessed. For the fear conditioning US contrast (startle white noise burst > implicit baseline), overlap between the auditory network and the activations was observed, as expected (**Figure 6)**. When assessed across all participants, the auditory network average activations were both higher than random (*T*(7) = 6.07, *p* < 0.001) and higher than the template auditory network (*T*(7) = 3.9, *p* < 0.01; **Figure 7)**. This was true even when examining auditory networks derived from rest only, although effects were smaller (vs. random, *T*(7) = 5.64, *p* < 0.001; vs template, *T*(7) = 3.02, *p* = 0.02; **Figure S21**). The only contrast to show similar overlap in Attention Network Task was Orienting (spatial vs center cue), wherein the frontoparietal network showed higher than random (*T*(7) = 3.45, *p* = 0.01) and higher than group (*T*(7) =3.20, *p* = 0.015). However, the overlaps were not significant when deriving the network from rest. No other contrast or networks showed significant overlap.

**Figure 6.**
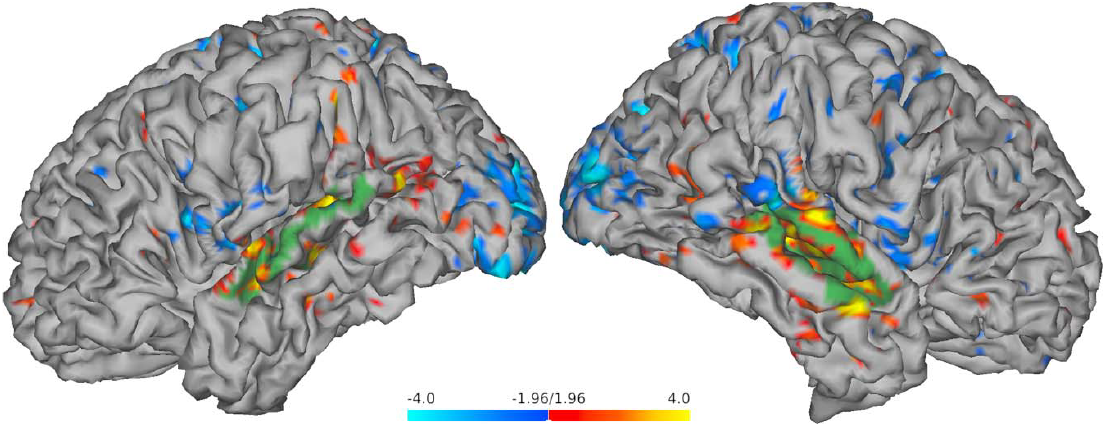
Surface display of activations (red-blue) and individualized auditory network (green) for an fMRI contrast isolating an auditory unconditioned stimulus (white noise burst > implicit baseline) during fear conditioning. Exemplar participant shown (1006). The activations were thresholded at a *Z*-score of 1.96, and the auditory network was overlaid. Inflated surface is found in **Figure S23**.

**Figure 7.**
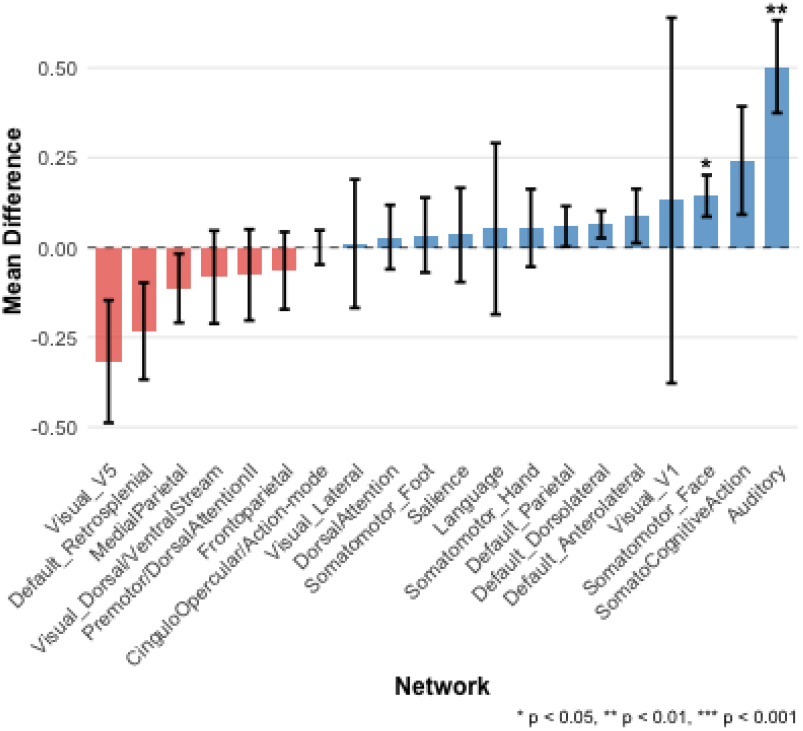
Individualized vs. group template differences in network activations during a fear conditioning task during presentation of an auditory unconditioned stimulus (white noise burst > implicit baseline). A positive difference implies that the individualized network had higher activations than the group network. Error bars reflect standard errors across participants.

## Discussion

In this preregistered case study, we investigated the feasibility and validity of PFM for identifying individualized brain networks in youth. We analyzed a dataset comprising ∼80 minutes of multi-echo data collected over one or two days, a duration that is long but feasible for developmental or other vulnerable populations. Analyses focused on the full ‘mixed-state’ data, including rest and task to maximize data quantity. In eight low-motion participants, we examined network topology, FC, and task-evoked responses. Overall, FC reliability compared favorably to prior single-echo studies, whereas stability estimates did not reach our preregistered thresholds. Sensory networks showed the highest spatial stability and task-optimized responses, whereas higher-order networks, such as the frontoparietal network and salience network, were less stable. Spatial stability was generally higher for the MS-HBM pipeline compared to Infomap, suggesting that different algorithms vary in sensitivity to FC differences. These results (**Table 2**) indicate that certain approaches may be more suitable than others for studying individualized network properties.

**Table 2.** Summary of Findings.

| **Individualized Network Property** | **Main Findings** | **Influence of Task Condition** |
| --- | --- | --- |
| <u>Network Topology</u> | Network sizes were not related to signal complexity or head motion. Sizes were largely similar to adult group templates. Pseudo-SE networks did not meet quality standards. | Sizes were similar across resting-state and mixed-state data, although salience network was larger in resting-state |
| <u>Network Topology Stability (Overall)</u> | Networks did not reach preregistered thresholds of stability, nor did stability reach an elbow point before 40 minutes. Algorithms affect overall levels of stability. | Network stability appears to be affected by task condition; resting-state stability increases faster than mixed-state |
| <u>Network Topology Stability (Specific Networks)</u> | Somatomotor, parietal and auditory networks show highest stability. Consistent across algorithms. DMN can reach high stability if ignoring subnetworks. | Consistent across task condition. |
| <u>Network Connectivity Reliability</u> | Connectivity reliability reaches an elbow point between 10-15 min of data. Overall reliability is highest for template networks, not individualized networks. | Connectivity reliability does not appear to be affected by task condition. |
| <u>Network Responsiveness to Tasks</u> | Individualized auditory networks showed optimized responses to auditory ‘shock’ stimulus from threat conditioning. | Optimized responses were noted even when defining networks on resting-state only. |

First, we examined the quality of the individualized networks. Networks showed modest variability and were similar in cortical extent to group template networks, with some differences in higher-order networks, like the default-mode and language networks. Although the purpose of the present work was not to examine developmental differences in network topology, it is notable that these findings are consistent with literature showing marked alterations in DMN and Language networks throughout development (F. Fan et al., 2021; Jasińska et al., 2021). Examinations of whole-brain FC demonstrated highly modular networks, with positive silhouette values across all identified networks. Modularity is a sought-after property of brain network parcellations because it indicates independence (Eickhoff et al., 2015). As such, there is evidence that multi-echo denoising and Infomap precision mapping results in good quality networks. The benefits of multi-echo have been previously established; in particular, multi-echo benefits SNR by allowing for theory-driven denoising of functional data (Kundu et al., 2017). Further, compared to conventional single-echo fMRI, multi-echo has been found to show higher stability of networks and reliability of FC in adults (Lynch et al., 2020).

To evaluate these possibilities in youth, we conducted rigorous and extensive tests of the stability of network topologies (spatial locations) over time. Our primary preregistered pipeline involved using Infomap, which finds communities in FC graphs by conducting random walks. Those communities are then assigned to networks (the quantity of communities depends on graph density thresholds). When applying Infomap, not only were stabilities increasing consistently over time (no elbows), but the overall overlap was below the 0.75 threshold previously applied in the adult literature (Gordon, Laumann, Gilmore, et al., 2017). Exceptions to this rule were found for specific networks, particularly sensory and perceptual networks. Specifically, the primary visual cortex, sensorimotor areas and superior temporal gyrus showed the most consistent network assignment.

Prior research has indicated that Infomap may be dependent on parameter choices (Laumann et al., 2015) and especially data-hungry (Hermosillo et al., 2024; Moore et al., 2024). Thus, we conducted an exploratory analysis of MS-HBM, which is a Bayesian prior method for network detection (Kong et al., 2019). MS-HBM is less computationally intensive than Infomap and resulted in more stable networks. Dice coefficients over 40 minutes of independent data increased more rapidly (20% to 26%), and a few participants showed elbows. Nevertheless, only one participant met the preregistered 0.75 threshold with 40 minutes of independent data. MS-HBM likewise showed highest stability in sensory networks (somatomotor, auditory, visual, and parietal). Interestingly, MS-HBM derivations of the auditory network showed similar task responsiveness to Infomap derivations, suggesting there were not significant individual sensitivity tradeoffs. While the current study was not designed to address algorithmic trade-offs in detail, our results suggest network topologies may continue to stabilize beyond 40 minutes, but that algorithms may influence results. These results are largely a replication of research in adults demonstrating gradual increases in stability and higher stabilities in sensory networks (Gordon, Laumann, Adeyemo, et al., 2017; Gratton et al., 2020; Naselaris et al., 2021).

On the other hand, when we examined the reliability of FC distributions across time increments, we observed high multivariate similarity within 10-15 minutes of independent data. Comparable reliability estimates have been reported in adults using multi-echo acquisitions (Lynch et al., 2020). Recently, a multi-echo study including adults, low-motion children, and high-motion children found that reaching high multivariate similarity took an average of 15.82 minutes in adults, 23.74 minutes in low-motion children, and 47.10 minutes in high-motion children (Rai et al., 2025). Further analyses established statistically that age and head motion have independent contributions to reliability. Single-echo reliability estimates in children are notably higher (45-50 minutes, Demeter et al., 2025). While other factors, such as analysis pipelines and viewing conditions may have effects, our results are generally consistent with prior work indicating multi-echo acquisitions may enable more reliable FC estimation than single-echo approaches, and suggest adolescents may approximate adult reliability (Lynch et al., 2020; Rai et al., 2025; Raut et al., 2019).

Finally, we examined task responses in the individualized networks. We found that the auditory network was the only network that showed consistently higher activations to a task than random and group networks, even when using held-out resting-state data to define the network. Although this may be related to the robust contrast (white noise burst > implicit baseline), this finding aligns with the broader sensory-first pattern: strong, time-locked sensory drives benefit most from individualized boundaries at feasible durations. Similar findings for sensory and motor networks have been observed in adults (Gordon, Laumann, Gilmore, et al., 2017; Laumann et al., 2015). Our findings that individualized auditory networks are especially task-responsive, and high in stability across independent data splits, suggest that PFM in limited quantities of adolescent multi-echo fMRI can capture individual, trait-like organization.

### Rest vs Task Considerations

In the current study, we chose to focus on mixed-state data to maximize data quantity. PFM has more commonly been performed using resting-state alone, but two studies have estimated individualized network maps applying the approach here, using mixed-state (concatenated rest + task) data without explicit task-design regression (Hermosillo et al., 2024; Wang et al., 2015). There is extensive debate over whether functional organization of brain systems (e.g., network shapes) differs across resting-state vs tasks (Cole et al., 2014; Gratton et al., 2018; Smith et al., 2009; Tavor et al., 2016). Thus, we conducted sensitivity tests to assess the impact of mixing states (**Table 2**). Results suggest that functional connectivity reliability may not differ between mixed-state vs. rest (reliability curves are similar, **Figure S19**), while there were mixed-state vs rest differences in network spatial stability (network stability grows more rapidly during rest only, **Figure S7)**. A previous study found low-demand conditions resulted in higher reliability, albeit with a small effect dwarfed by motion and age (Rai et al., 2025). It is possible that these results may differ with other pipelines like task-design regression (see Kraus et al., 2021), and future work should further examine whether task condition differences are most influential on functional connectivity reliability or network topology.

### Limitations

The current case study focused mostly on visual interpretation of reliability and stability curves. Many interpretations could be better supported by statistical analyses in large cohorts (e.g., does MS-HBM provide higher stability than Infomap? what are FC reliabilities for multi-echo and single-echo?). It should be mentioned that the current case study allowed computationally intensive precision mapping within-person across increasing time windows, which may not be possible for large samples.

Another concern regards FC reliability. We highlight our multivariate similarity results, as they are most comparable to previous PFM studies in youth (Demeter et al., 2025; Rai et al., 2025). Multivariate similarity captures the reliability of the large-scale organization of the brain. However, multivariate similarity depends on the underlying features (e.g., parcels, networks, or vertices), and we found a pattern where reliability was highest for group networks > individualized networks > vertices. One possibility is that individualized networks did not improve FC reliability here because network organization itself may have been limited in stability.

Further, patterns were meaningfully different when examining a different operationalization of reliability, edge-wise ICC, which explicitly models the ratio of between-individual variance to within-individual variance. While our elbow analysis also identified a point of diminishing returns at 10-15 minutes in ICCs, this is not an asymptote and there were continued increases after the elbow point (increasing from 0.43 – 0.57 from 10-40 minutes). This is consistent with previous literature noting modest univariate ICCs for many networks even when multivariate similarity is high (Rai et al., 2025). Importantly, despite these modest ICC values, reliability estimates in the current multi-echo dataset appear comparatively favorable relative to prior single-echo studies. For example, network-level ICCs observed here with approximately 10 minutes of multi-echo data are comparable to or higher than some single-echo estimates reported using substantially longer acquisitions (>30 minutes) (Noble et al., 2017, 2019). Nevertheless, reliability requirements may differ depending on the intended application (e.g., multivariate pattern estimation vs. individual edge estimation, or within-individual vs. between-individual inference), and future work should continue evaluating these distinctions.

Additionally, reliability and stability should be considered in the context of experimental questions. While high reliability fMRI measures are often sought out (Noble et al., 2017), reliability in a vacuum is not always positive. Hair color may be extremely reliable in adolescents, but not predictive of any cognitive traits. As such, comparatively better or worse reliability in our analyses may be associated with other differences (e.g. individual specificity) that may be more relevant for a given study application. Our current work suggests that multi-echo data may provide high-quality fMRI data not only because of its reliability, but also because the networks show other properties like meaningful deviations from templates, sensitivity to tasks, and lack of sensitivity to structural confounds.

### Recommendations

Based on these findings, we offer three recommendations for future studies of individualized networks in adolescents. First, multi-echo acquisition may improve the reliability of FC estimates—particularly multivariate metrics—consistent with its theory-driven denoising framework. Second, caution is warranted when interpreting PFM-derived networks as trait-like individual differences. In our data, overall spatial stability did not reach preregistered thresholds or elbow points even after 40 minutes of multi-echo acquisition, although sensory networks were comparatively stable and task-responsive. Third, network stability may vary by task condition, suggesting that studies combining task and rest data to increase sample size should adopt principled strategies to account for task-related effects. These recommendations should be interpreted cautiously, given ongoing challenges in generalizing findings across fMRI preprocessing pipelines (Botvinik-Nezer et al., 2020) and across samples (Ellwood-Lowe et al., 2021; Greene et al., 2022).

## Conclusions and Future Directions

In summary, our case study reveals the promise and limitations of adolescent multi-echo fMRI for investigating individualized brain functional organization. Our findings indicate that with modest scan times, multi-echo acquisition may improve FC reliability relative to prior single-echo estimates. By contrast, with limited scan time, stable individualized network topology emerges most readily in sensory systems and remains constrained in association cortex. Moreover, stability depends on PFM algorithms, and state (task condition) effects may be identified. Our findings encourage expanded use of multi-echo PFM in youth, while underscoring the need for careful navigation of these tradeoffs and further methodological innovation.

### Ethics approval

All procedures complied with the Declaration of Helsinki and were approved by the Wayne State University Institutional Review Board. A parent or legal guardian provided written informed consent and all youth provided informed assent.

### Sex- and gender-based analysis

Sex (assigned at birth) was recorded. Given the small, feasibility sample, sex-/gender-based analyses were not performed.

## Data and Code Availability

De-identified data and code will be made available upon acceptance at https://osf.io/kgvsb/overview?view_only=73be63e38b18407da85a6bb85bd38602.

## Author Contributions

**Conceptualization:** Treves

**Methodology:** Treves, Pagliaccio, Patel

**Software:** Treves

**Validation:** Treves, Pagliaccio

**Formal Analysis:** Treves

**Investigation:** Treves, Tamimi, Marusak

**Writing - Original Draft:** Treves

**Writing - Review & Editing:** Treves, Pagliaccio, Tamimi, Patel, Kimerty, Auerbach, Marusak

**Visualization:** Treves, Pagliaccio, Kimerty

**Supervision:** Auerbach, Marusak

**Project administration:** Tamimi, Marusak

**Funding acquisition:** Marusak

## Acknowledgements

The authors would like to thank the following individuals and organizations for their assistance with recruitment, data collection, and/or analysis: Amanpreet Bhogal, Shelley Paulisin, Jovan Jande, Emily Crisan, Alexander Jakubiec, Shravya Chanamolu, Sneha Bhargava, Myles Davis, Zazai Owens, Carla Hannah, Iveta Kopil, Shreya Desai, Cameron Martella, Autumm Heeter, Julia Evanski, Ahmad Almaat, Leah Gowatch, Dr. Laura Benjamins, Dr. Sharon Marshall, Dr. Christopher Youngman, Dr. Clara Zundel, Samantha Ely, Carmen Carpenter, MacKenna Sahmpine, Reem Tamimi, Mariya Matsko, Sarah Rogers, Jennifer Losiowski, Emilie O’Mara, Kamakashi Sharma, Wayne Pediatrics, The Children’s Center, The Wayne State University MR Research Facility.

## Declaration of AI use

AI tools (CLAUDE SONNET 4) were applied for refining MATLAB code and R code for analysis and visualizations. After using this tool, the author(s) reviewed and edited the content as needed and take(s) full responsibility for the content of the published article.

