## Supplement for "Feasibility of Precision Functional Mapping in Youth Multi-Echo fMRI Data"

**Supplement Text 1: Scan Parameters**

Functional MRI data were acquired using a multi-echo gradient-echo echo-planar imaging (ME-EPI) sequence with the following parameters: repetition time (TR) = 1500 ms; echo times (TEs) = 13.6, 30.87, 48.14 ms; flip angle (FA) = 83°; 51 axial slices; field of view (FOV) = 209 × 209 mm²; voxel size = 2.9 mm isotropic; parallel imaging acceleration factor = 2. Images were acquired with no gap between slices, providing whole-brain coverage.

**Supplement Text 2: Scan Order**

Day 1 - resting state 1, fear conditioning, resting state 2, extinction, resting state 3.

Day 2 - ANT, resting state 4, recall, renewal.

Scan order was randomized for each individual for independent splits.

**Supplement Text 3: Participant Selection**

Out of 80 possible participants, 8 were selected based on right-handedness, low head motion, adequate task performance, and a lack of scan notes. The head motion threshold was < 25 frames scrubbed out of a total of 3200 frames (based on threshold of 0.3 mm). This resulted in mean framewise displacements between 0.033 and 0.066 mm. For the tasks, the goal was to assure compliance – they were required to have a high rate of response during Fear Conditioning, and a high accuracy during the Attention Network Task. Scan notes for resting-state consisted of noting whether they fell asleep (no case study participants fell asleep).

**Supplement Text 4: Fieldmap-less correction**

Fieldmaps were not collected based on the prior assumption that multi-echo denoising could account for distortions. However, in our PFM examinations, fieldmaps were found to make a difference for quality. Given the developmental population, instead of using an adult fieldmap template, we averaged fieldmaps from a separate sample of 100 adolescents previously reported in Treves et al., 2024.

**Supplement Text 5: MS-HBM**

MS-HBM, i.e. multi-session hierarchical Bayesian model (Kong et al., 2019), was a computationally inexpensive method used to derive individualized networks. This method involves comparing (via likelihoods) vertex-level connectivity maps to weighted group priors to derive network assignments. Prior work has demonstrated moderate spatial overlap (NMI = 0.63) between networks derived from MS-HBM and Infomap (Lynch et al., 2024) Two free parameters govern the network identification – the smoothness across vertices, and the prior weighting. Other parameters – inter-subject variability in connectivity profiles, intra-subject variability across sessions, and group priors – are pre-set in the algorithm and learned from a large training set.

**Supplement Text 6: Case 1003 Outlier**

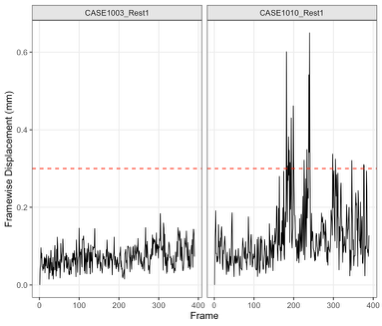

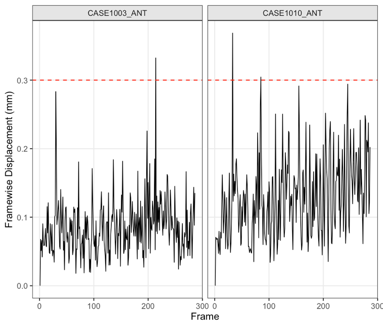
Case 1003 demonstrated high FC similarity after only 2.5 minutes of independent data (0.9). Notably, Case 1003 showed lower motion than the other subjects, with only 4 timepoints removed (FD threshold > 0.3 mm) and an average of 0.033 mm FD. Example FD traces are shown below for 1003 and 1010 for example runs. There is no evidence of performance differences in this subject, and the subject was not asleep.

**Supplement Text 7:** **Pseudo-SE Results in Sub-Optimal Networks**

For a pseudo single-echo comparison, the middle-echo images were used (Lynch et al., 2020), and then preprocessed identically. Instead of *tedana* denoising, the ICA-AROMA denoising method was applied (Lynch et al., 2024; Pruim et al., 2015). As shown in Figures S4 and S5, both Infomap and MS-HBM resulted in poor network quality using the middle-echo. Thus, it is likely caused by the denoising pipeline. While we followed one published approach (Lynch et al., 2024), other groups have used different approaches incorporating filtering, noise component regression, and multiple detrending and demeaning steps(Chernicky et al., 2026; Gratton et al., 2020; Kucyi et al., 2024). Interestingly, the multi-echo pipeline incorporated a low-threshold template similarity step to remove components not matching group networks, which was not possible to implement in ICA-AROMA. Different denoising steps may lead to meaningful differences in vertex-level connectivity, in turn impacting the quality of networks derived using Infomap or MS-HBM.

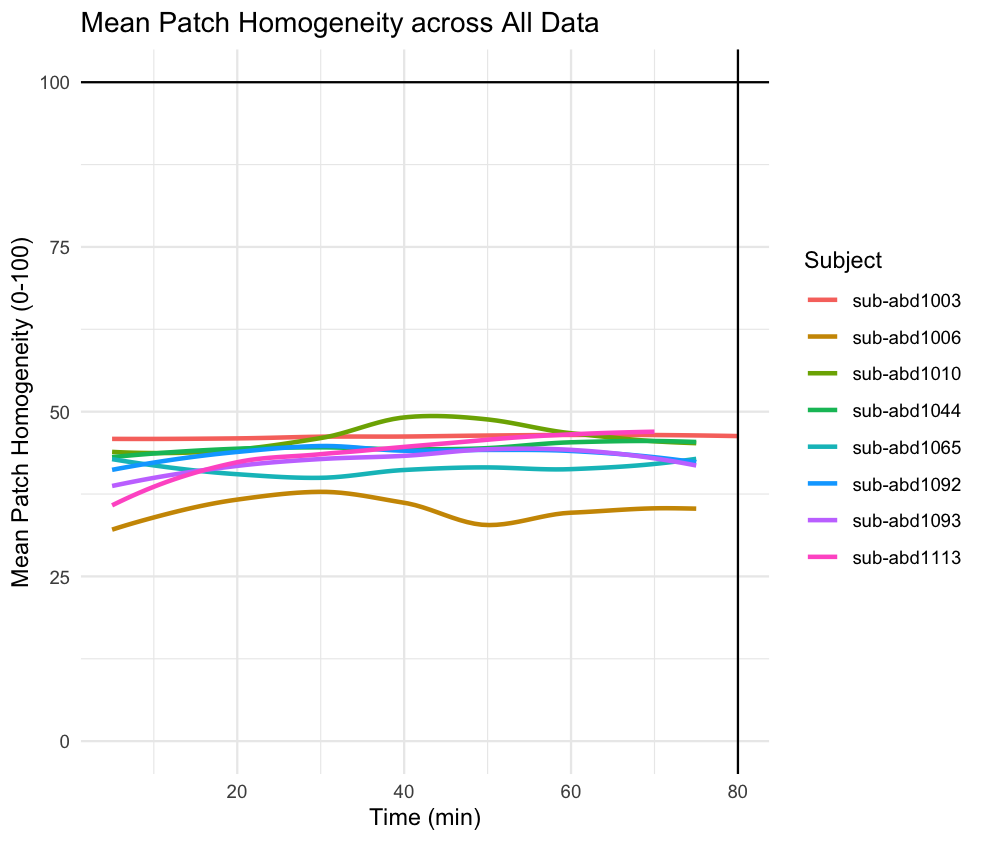

**Figure S1:** Homogeneities (percentage variance explained) over time for full data for Infomap pipeline.

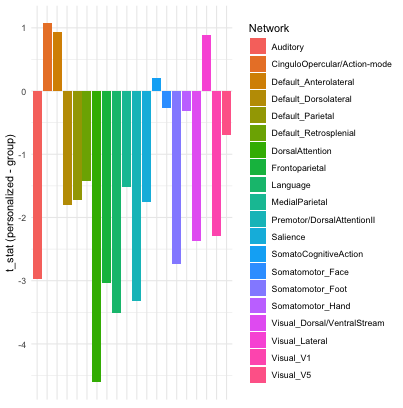

**Figure S2:** Homogeneities of each network from Infomap compared to homogeneities of group networks. A negative t-stat reflects more homogeneous activations in the group network. Group networks show consistently larger timecourse homogeneity than individualized networks.

**B**

**A**

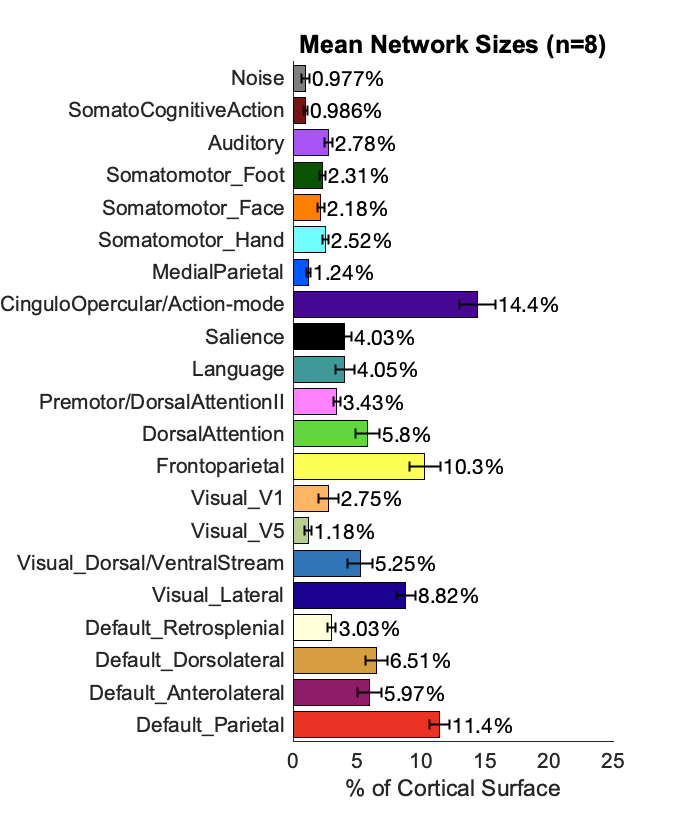

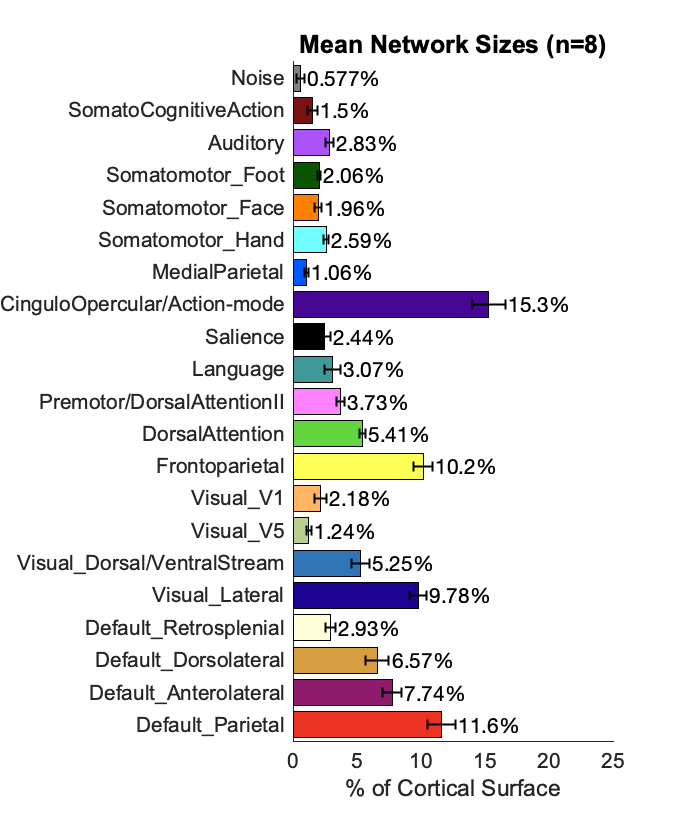

**Figure S3: Individualized network surface areas for full data (A) vs rest (B).** Error bars reflect standard errors across participants. Particularly notable is that the salience network in the full data occupies an average of 2.44% of the cortical surface, and in rest data it occupies an average of 4.03% of the cortical surface, a substantial increase.

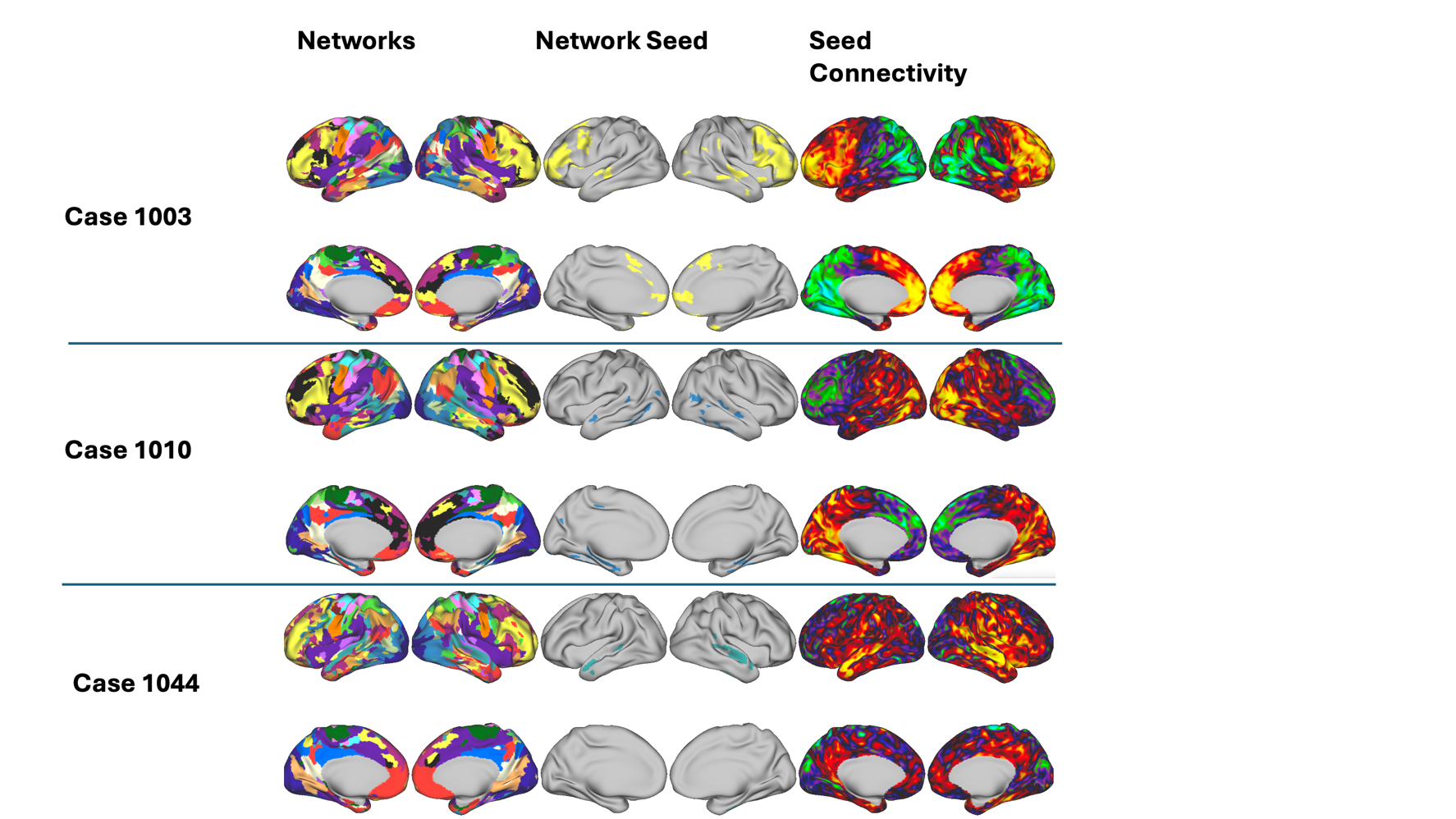

**Figure S4: Infomap network maps for pseudo-SE data.** For subject 1003, note the abnormal positioning of the frontoparietal (yellow) and retrosplenial cortex (white), as well as gradient connectivity. For 1010, note the massive salience network, and gradients. For 1044, note the massive medial parietal network and noisy connectivity (although large-scale network connectivity patterns are evident). For discussion of sources of pseudo-SE problems, see **Supplement Text S7**.

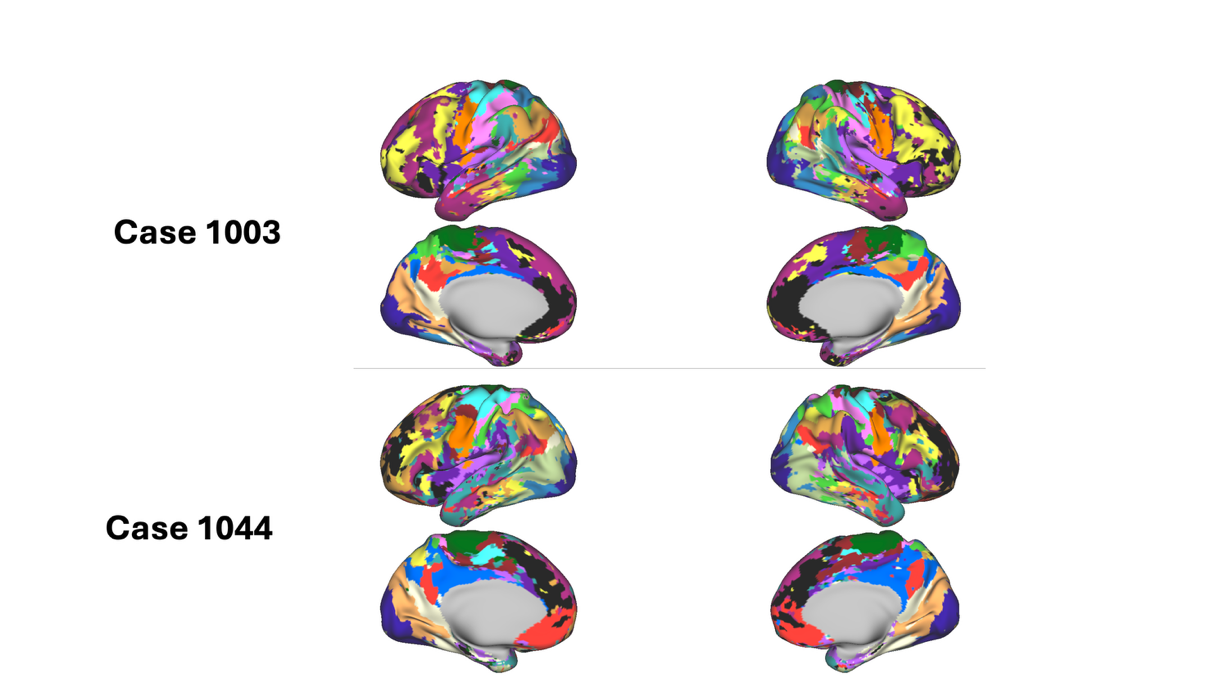

**Figure S5: MS-HBM network maps for pseudo-SE data.** There are clear problems with the network maps, including overrepresentation of the salience network (black) and cingular opercular network (purple). These problems are found with Infomap as well, suggesting the raw timecourses/FC used to generate the maps may contain diffuse noise. For discussion of sources of pseudo-SE problems, see **Supplement Text S7**.

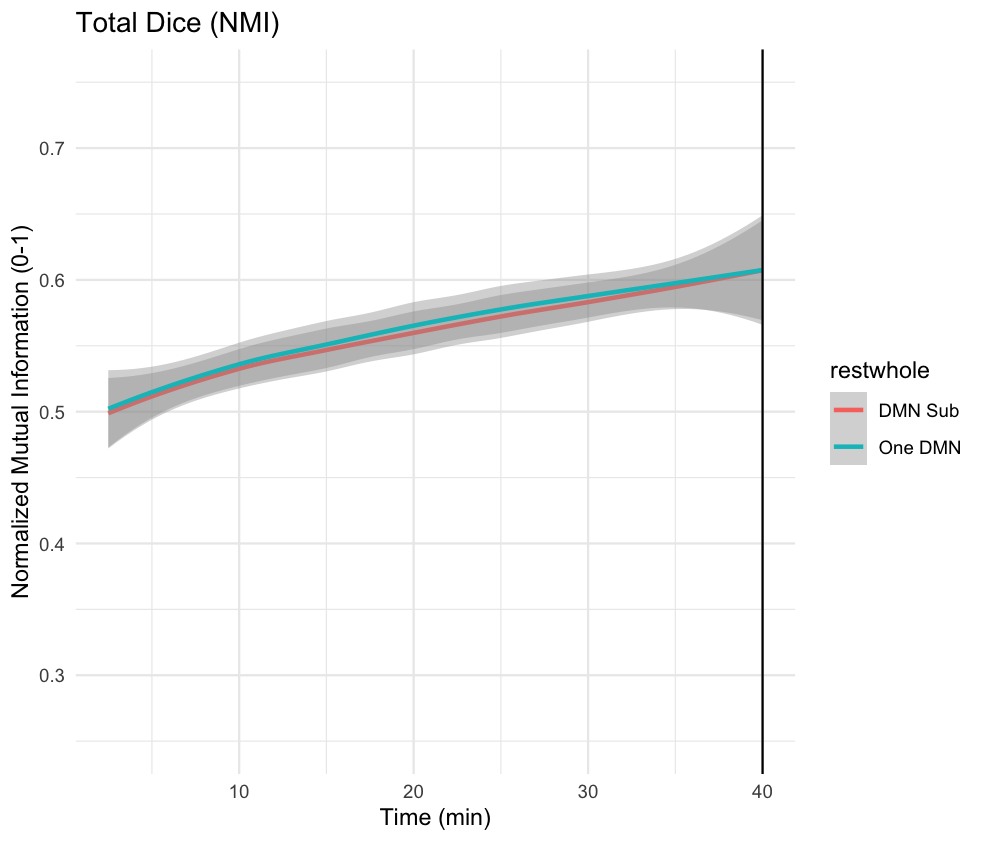

**Figure S6:** Overall network stability as measured by normalized mutual information (similar to dice), when combining DMN networks (‘One DMN’) or examining subnetworks (‘DMN Sub’).

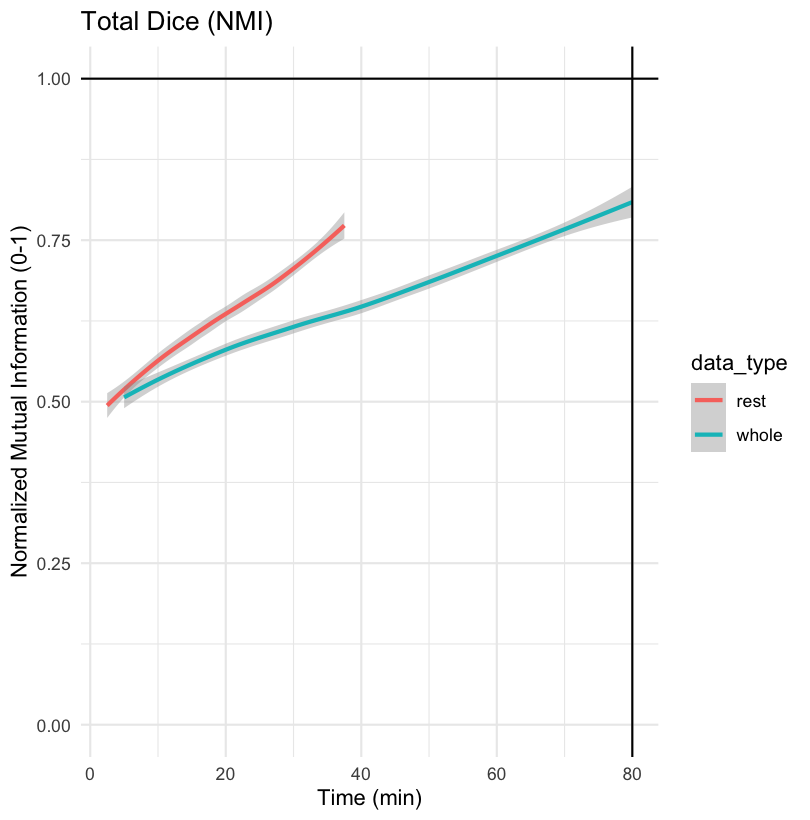

**Figure S7:** Overall network stability for Infomap pipeline comparing resting-state and mixed-state (whole). Normalized mutual information comparing the similarity of network topology across all networks over time increments of data. Trajectories of stability are shown in minutes. In blue, all data is shown, consisting of up to 80 minutes. In red, resting state data is shown, consisting of up to 37.5 minutes. A more rapid increase is observed in resting-state only data, suggesting possible state effects.

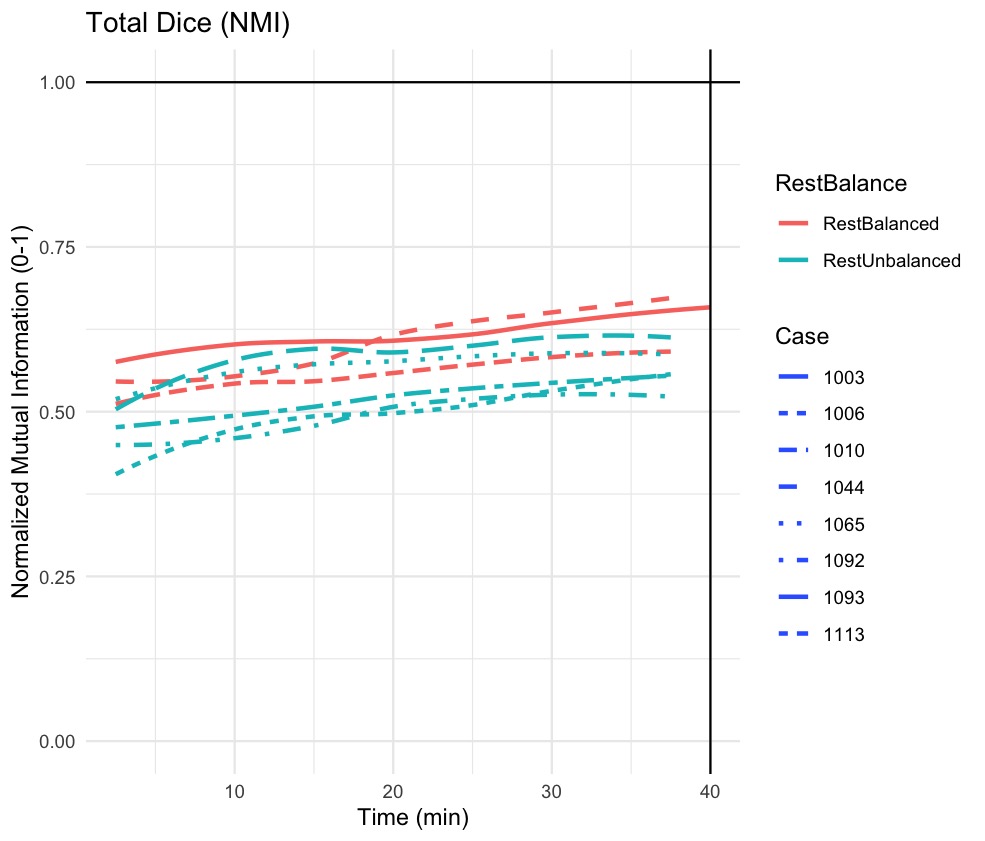

**Figure S8:** Dice stability (Infomap) curves for each participant. Participants were separated into ‘RestBalanced’ and ‘RestUnbalanced’. Participants with ‘RestBalanced’ had equal amounts of resting-data in the training data and held-out data. Here, dice stability seems to be higher when rest data is balanced, although the small subject pool limits this inference.

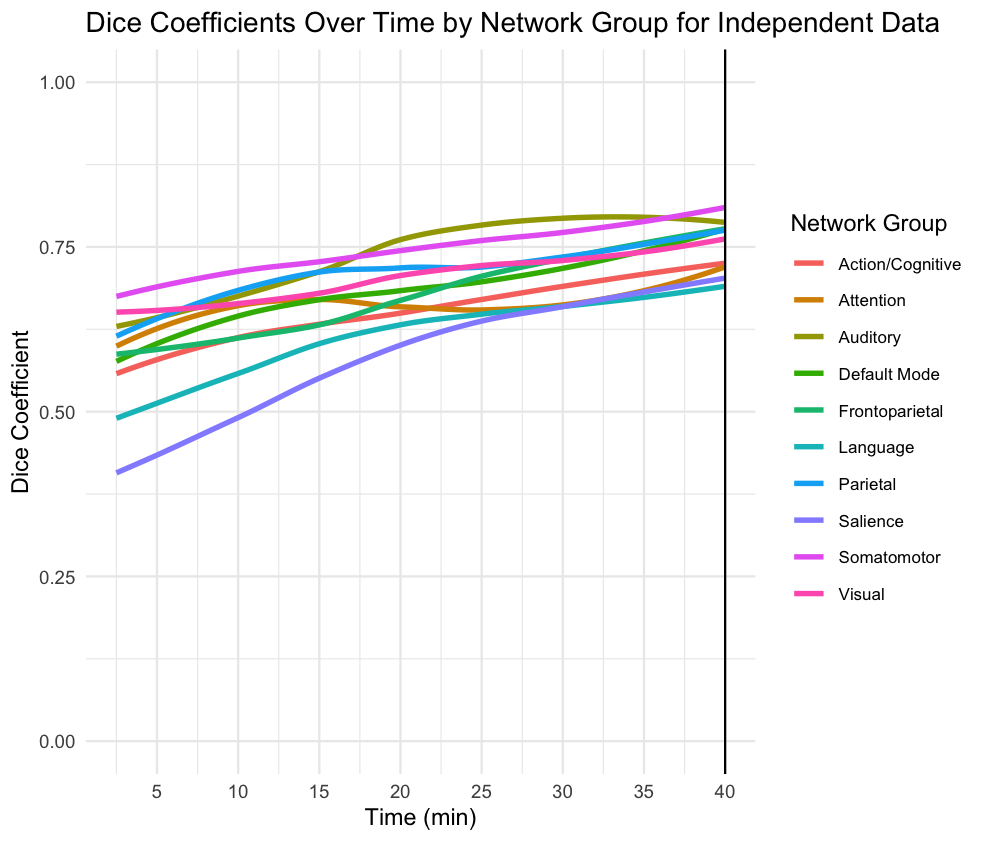

**Figure S9:** Dice coefficients by network over increasing amounts of data, with MS-HBM pipeline.

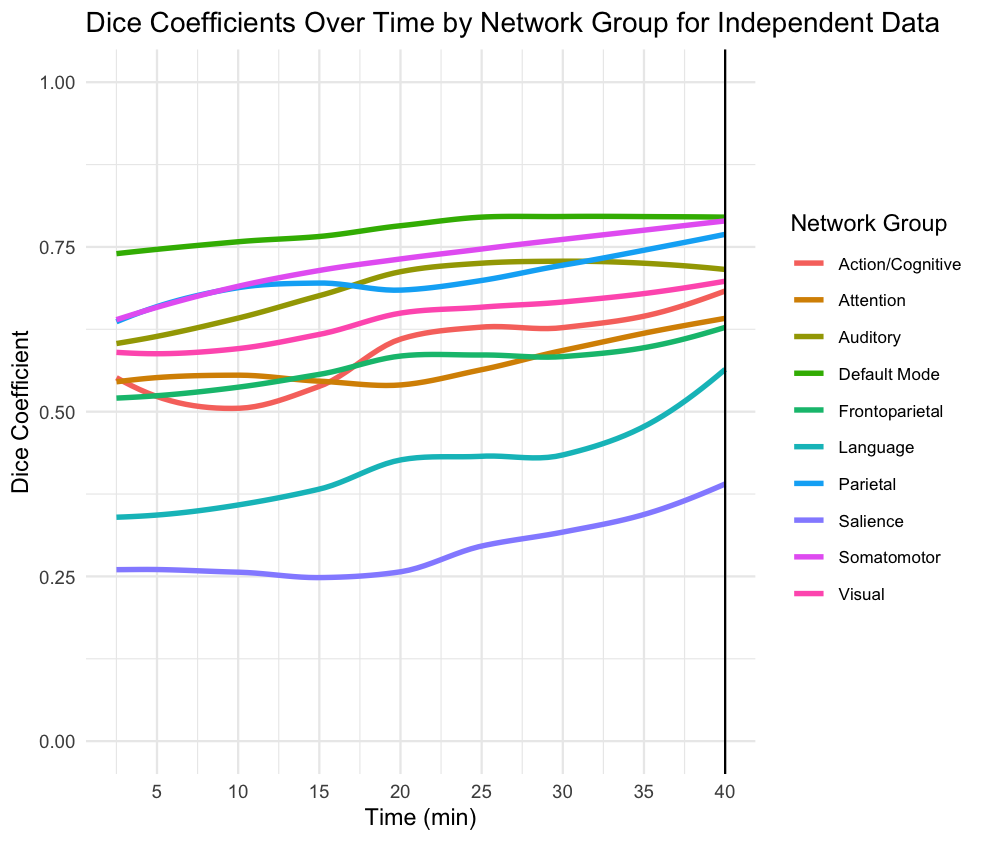

**Figure S10:** Dice coefficients by network for Infomap pipeline, with DMN networks combined before calculating coefficients (green).

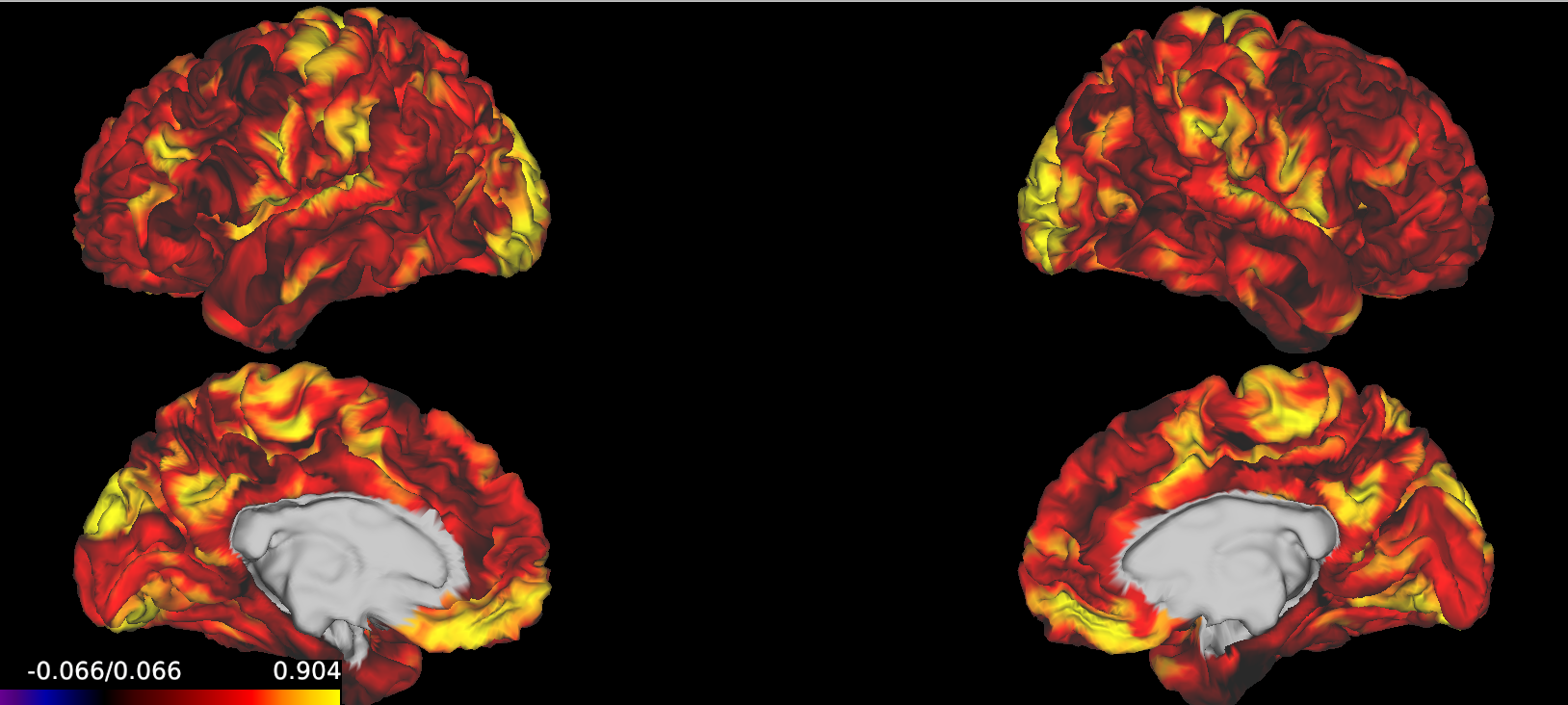

**Figure S11:** Average proportion of splits assigned to group template network vs distinct network. Yellow is more frequent. Red-black is less frequent.

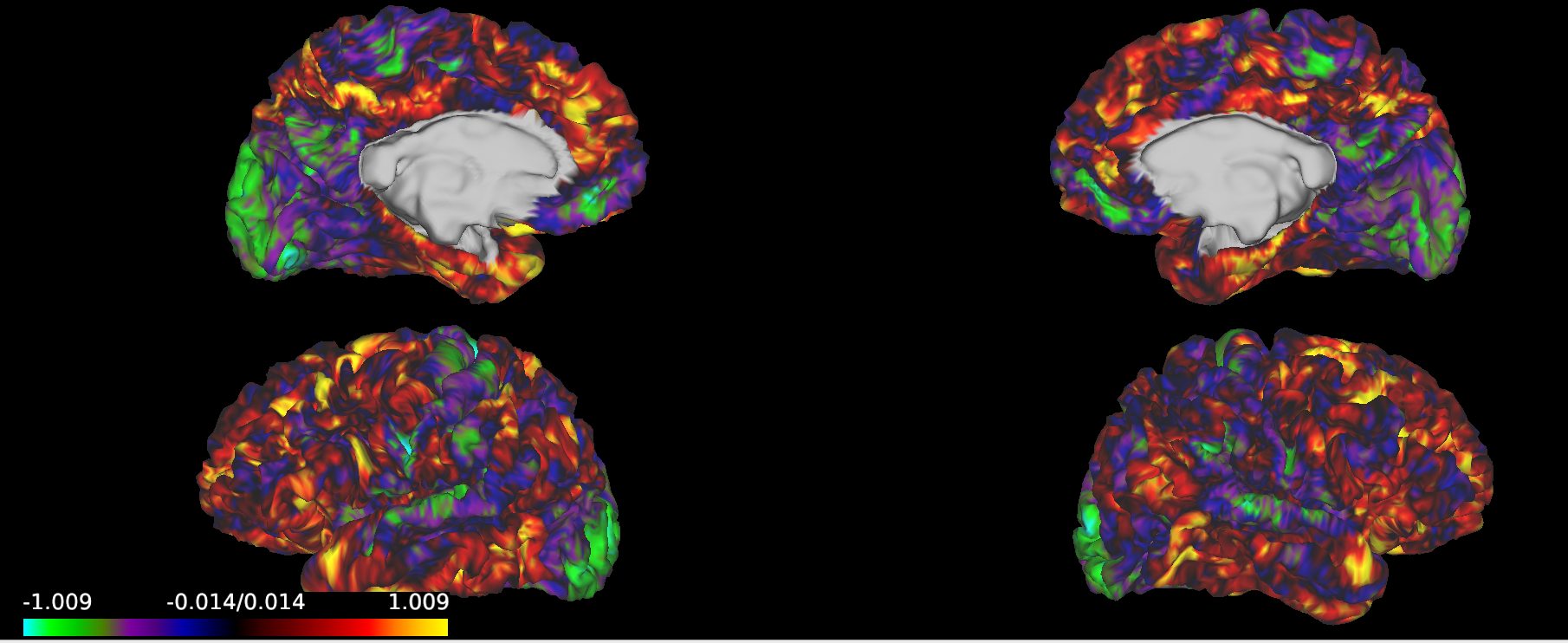

**Figure S12:** Number of unique networks (z-scored) for each brain vertex. Green reflects few networks , yellow reflects many networks.

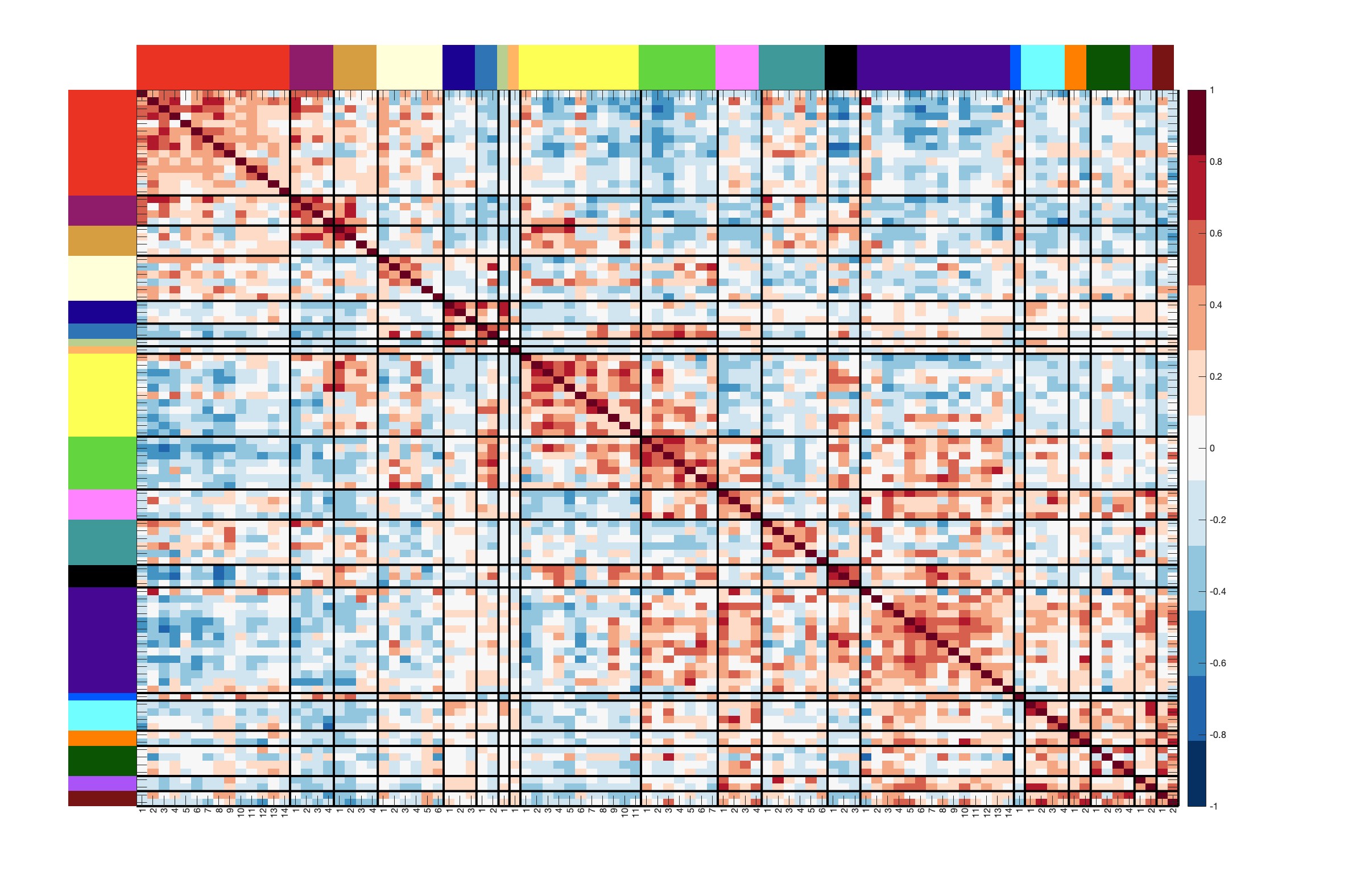

**Figure S13:** Functional connectivity for sub-abd1006. Colors represent distinct networks, see Figure S1. Red connections reflect high positive connectivity, and blue connections reflect low negative connectivity.

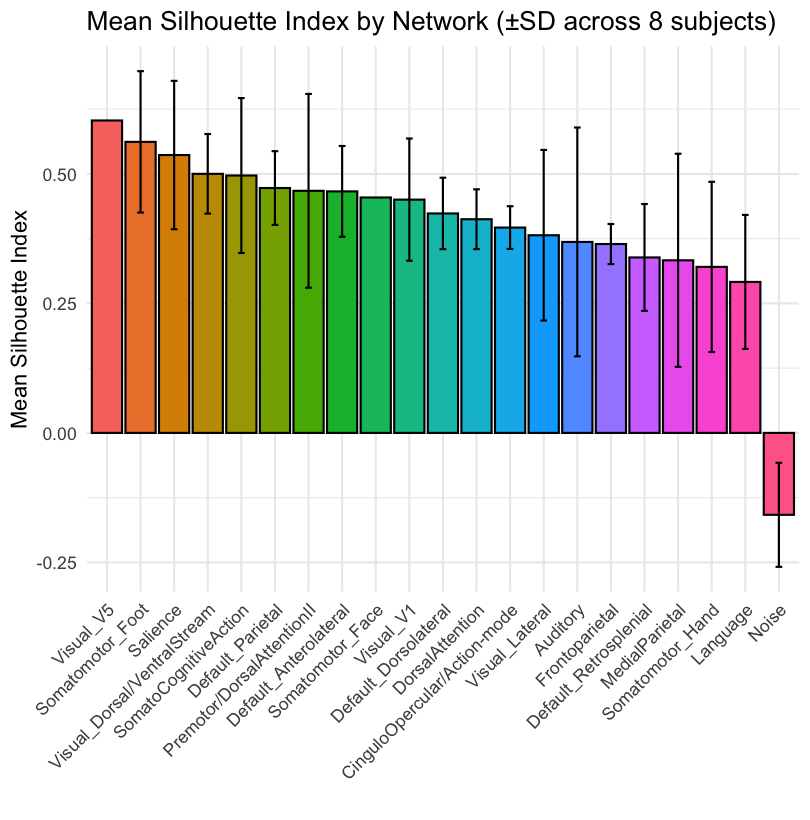

**Figure S14**: Silhouette values for individualized networks, defined as (within-network connectivity – between-network connectivity) / (within_network connectivity +between_network connectivity). Error bars (SD) across individuals are shown. Note that for all but two individuals (1044/1093), Visual_V5 and Somatomotor Face networks typically consisted of single communities, as such the Silhouette Index was not defined. 1044 had a large and positive silhouette value for Visual_V5, and 1093 had a large positive value for Somatomotor Face; there are no error bars given only one participant. The Noise network was a category for networks and connectivity that could not be categorized otherwise, showing negative silhouette values.

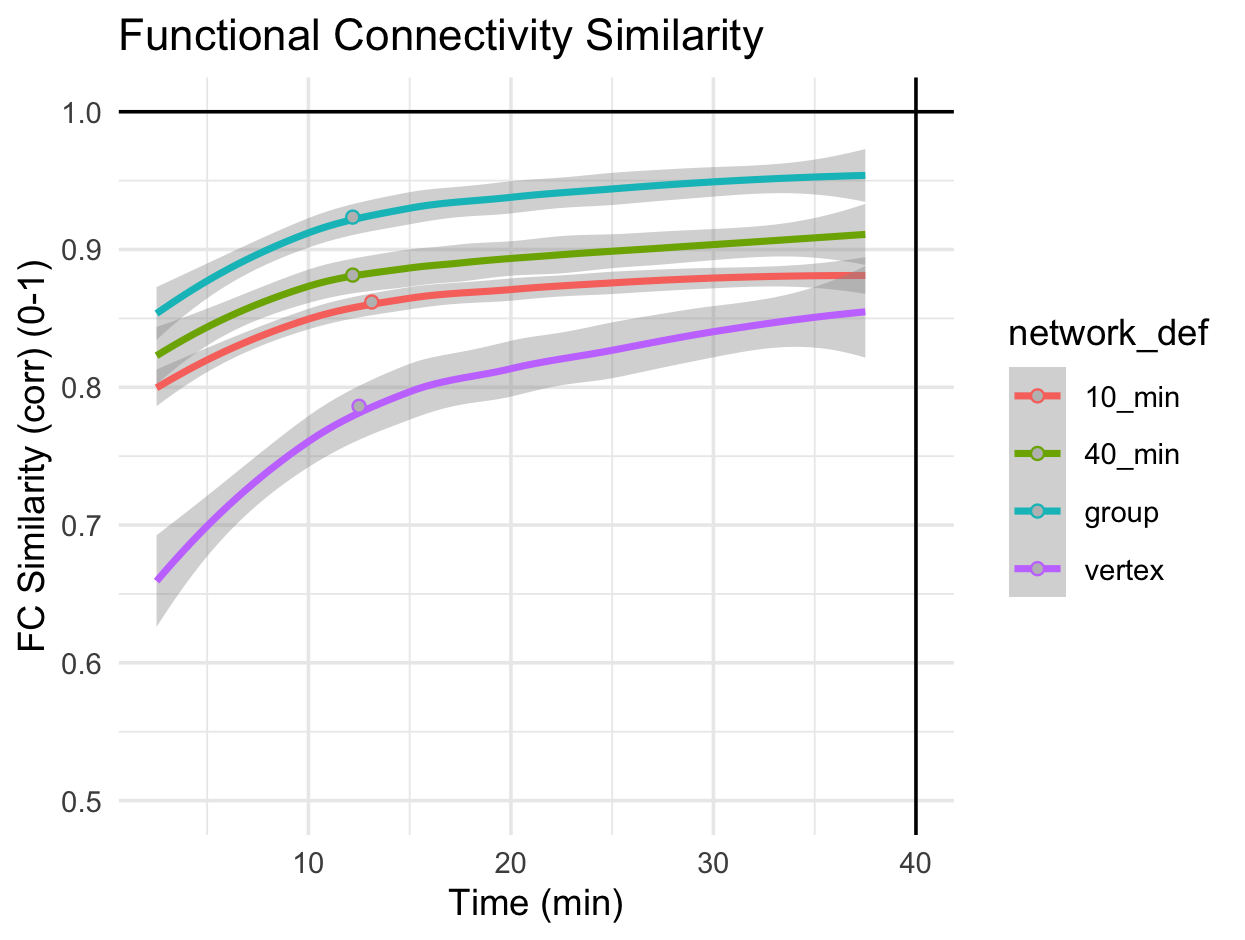

**Figure S15:** FC reliability in independent data. Group: networks from template. 40 minutes: networks defined on entire training data. 10 minutes: networks defined on 10 minutes of training data. Vertex: average FC similarity of vertex connectivity, excluding close vertices (Lynch et al., 2020). The mean elbow (across participants) of each curve using the Unit Invariant Knee (UIK) method is noted by a gray point.

For 10_min, the mean elbow was 13.1 min (range=10-20) yielding similarity of mean=0.862 (range 0.818-0.895). For 40_min, the mean elbow was 12.2 min (range=10-15) yielding similarity of mean=0.881 (range 0.836-0.964). For group, the mean elbow was 12.2 min (range=10-15) yielding similarity of mean=0.924 (range 0.851-0.962). For vertex, the mean elbow was 12.5 min (range=5-17.5) yielding similarity of mean=0.786 (range 0.599-0.870).

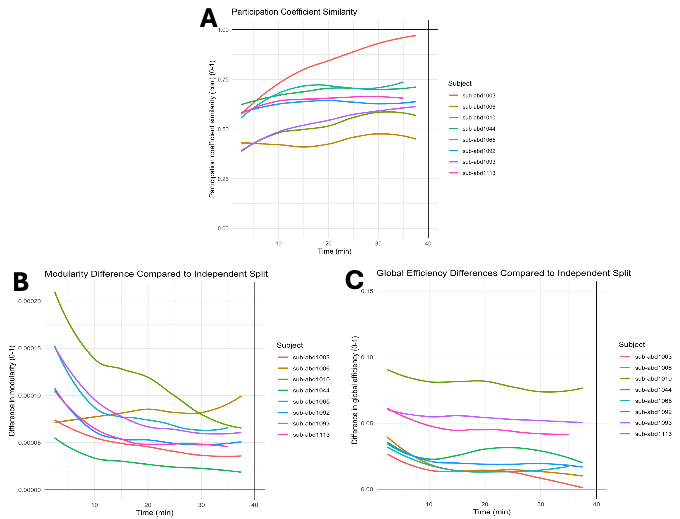

**C**

**Figure S16:** Differences in graph theoretic properties of individualized networks with increasing time. A) Participation coefficient represents the similarity of participation coefficients for each network for each split of the data. For B) modularity and C) global efficiency, values were calculated across each split and then the absolute value of the difference is plotted. Participation coefficient (PC) is a measure of the degree to which a graph node is connected to multiple networks. Modularity is a whole-graph measure representing the degree to which the graph can be well represented as a set of discrete modules. Global efficiency is a whole-graph measure of how efficiently information can be transferred over a graph. See (Gordon et al., 2017) for full definitions.

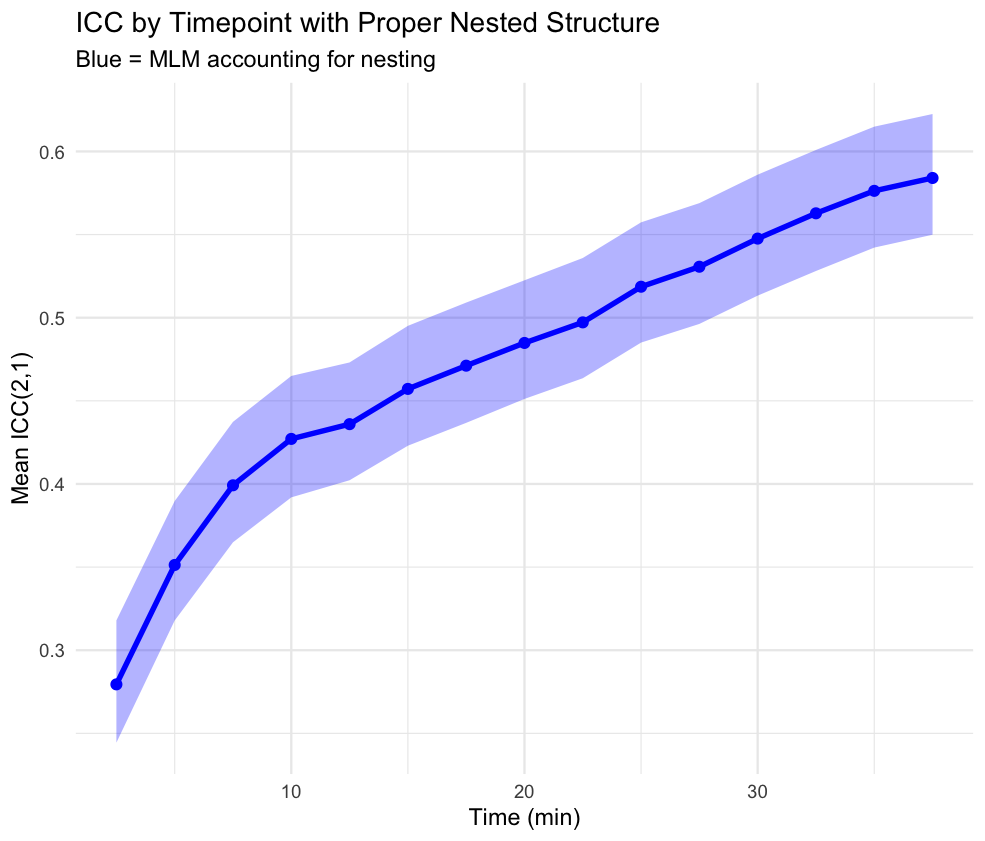

**Figure S17:** Mean trajectories of network-network ICC in independent data, employing individualized networks. Network-network connectivity was calculated across shuffles of training data (e.g., random 10 minute partitions) and then compared to the ground truth held-out data using ICC(2,1), and then averaged across individuals. Blue shading reflects multilevel model confidence intervals. Higher ICC reflects more between-person variance than within-person variance. Like FC similarity, ICC trajectories showed an elbow around 10 minutes. Like FC similarity, ICC trajectories showed an elbow at 10 minutes (gray point - using the Unit Invariant Knee (UIK) method). However, unlike FC similarity, ICC values continued to increase steadily through 40 minutes of data.

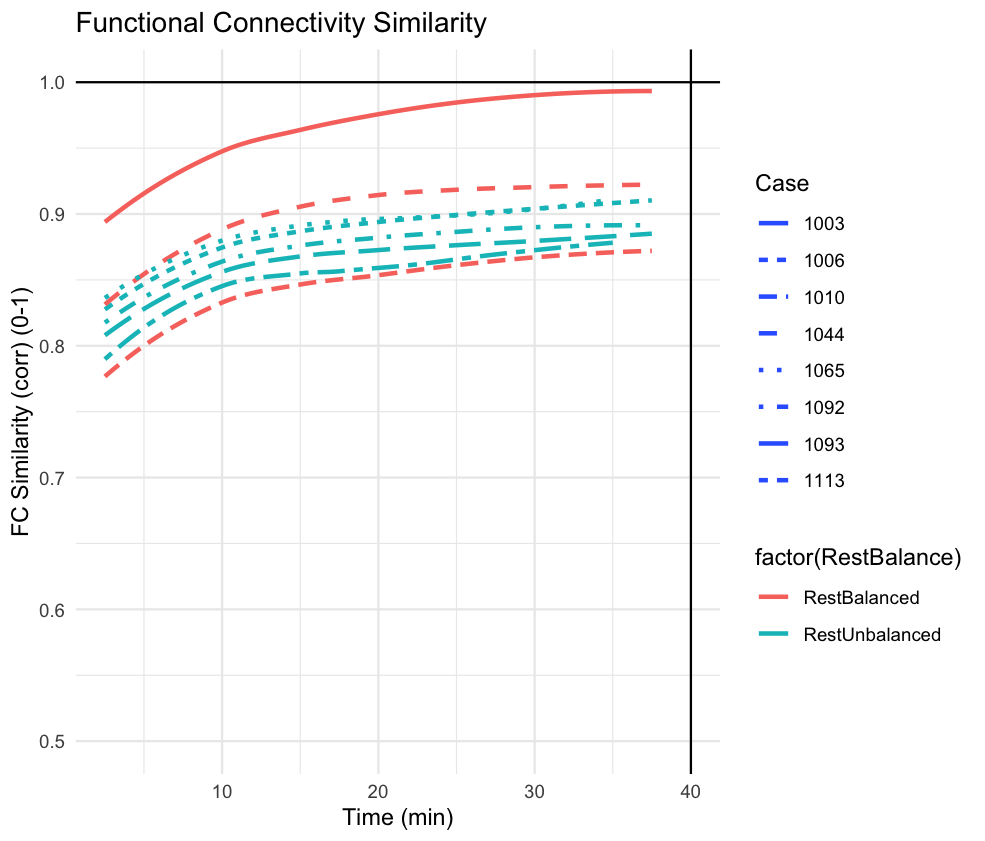

**Figure S18:** FC similarity curves for each participant. Participants were separated into ‘RestBalanced’ and ‘RestUnbalanced’. Participants with ‘RestBalanced’ had equal amounts of resting-data in the training data and held-out data. Here, FC similarity does not appear to be significantly different depending on rest data similarity, although the small subject pool limits this inference.

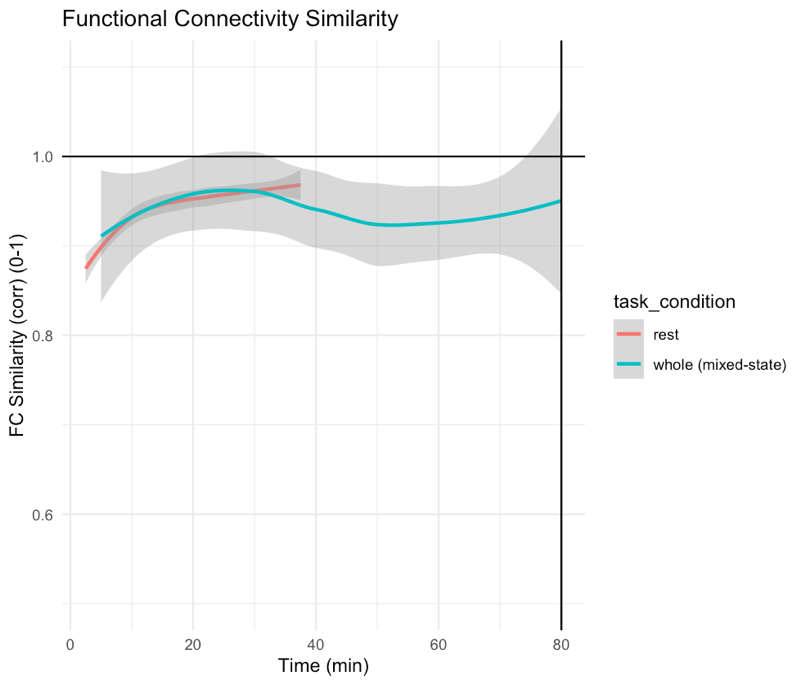

**Figure S19:** FC similarity curves for non-independent data splits. Trajectories of FC similarity are shown in minutes. In blue, all data is shown, consisting of up to 80 minutes (starting at 5 minutes). In red, resting state data is shown, consisting of up to 37.5 minutes (starting at 2.5 minutes). Standard errors were not restricted to (0-1) for ease of display, although FC similarity above 1 is not possible. Trajectories appear similar.

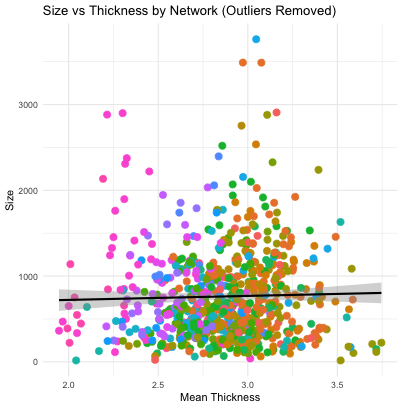

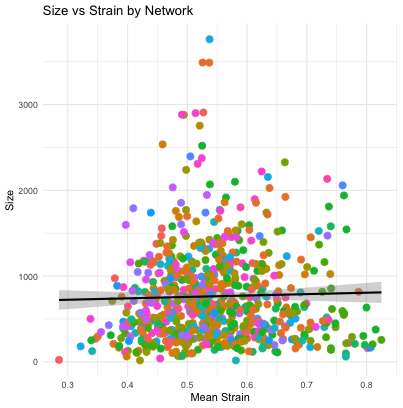

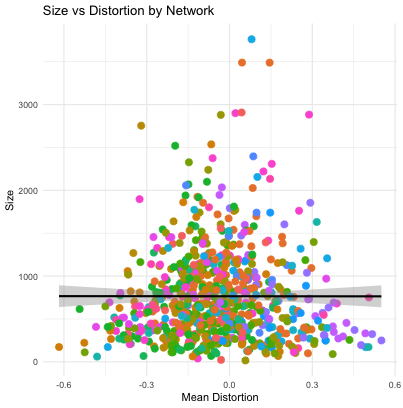

**Figure S20:** Structural metrics vs network surface area (size). Left, cortical thickness vs surface area. Middle, rotational strain vs surface area. Right, rotational distortion (expansion /contraction) vs surface area. Colors reflect distinct networks. Black line reflects best linear fit , with gray standard error bars. No significant relationships were found (*ps* > 0.25).

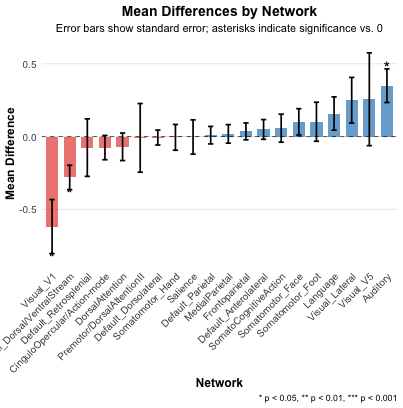

**Figure S21:** Resting-state defined networks and activations during shock. Auditory cortex shows significantly larger activations than group network. Ps are uncorrected for multiple comparisons based on the extremely robust hypothesis that auditory cortex will respond to a surprising auditory stimulus.

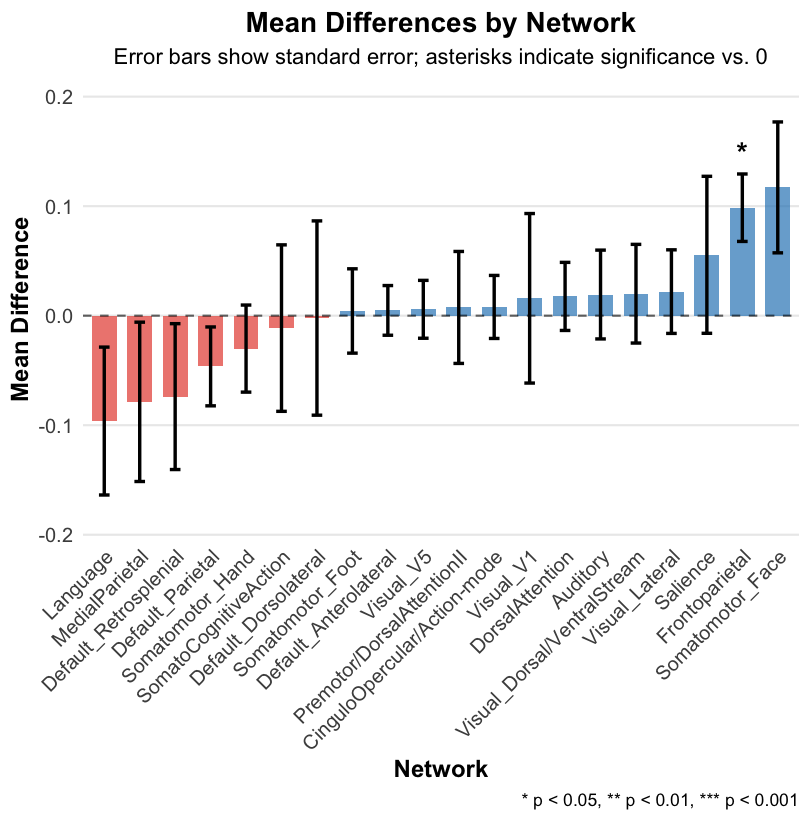

**Figure S22:** Full data defined networks and activations during orienting. Frontoparietal network shows significantly larger activations than group network. Ps are uncorrected for multiple comparisons, based on hypothesis that frontoparietal network is involved in attention orienting.

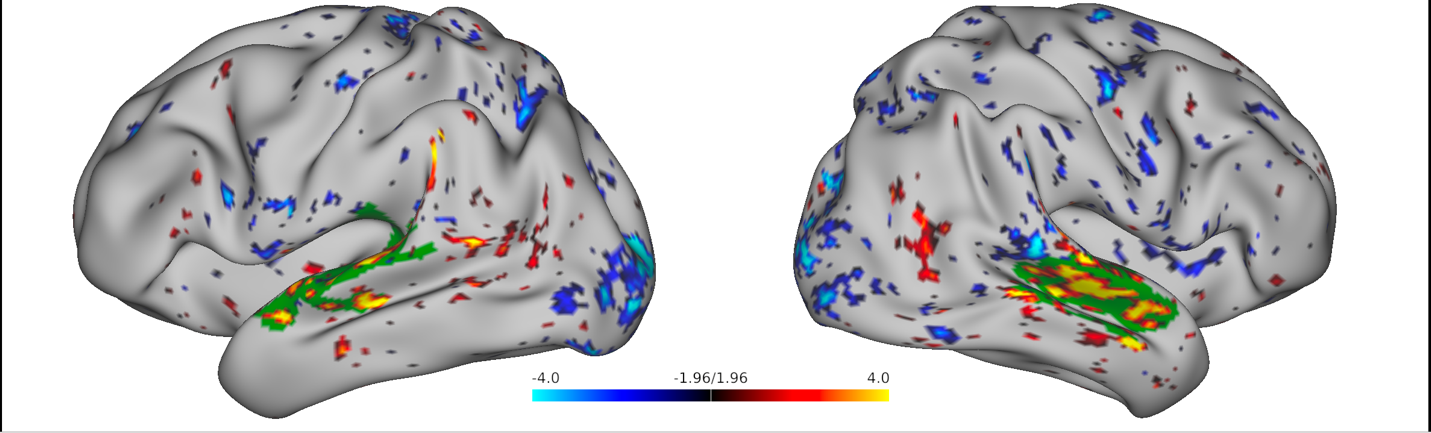

**Figure S23:** Surface display of activations (red-blue) and individualized auditory network (green) for an fMRI contrast isolating an auditory unconditioned stimulus (white noise burst > implicit baseline) during fear conditioning. Exemplar participant shown (1006). The activations were thresholded at a *Z*-score of 1.96, and the auditory network was overlaid.
